# Active prokaryotic and eukaryotic viral ecology across spatial scale in a deep-sea brine pool

**DOI:** 10.1101/2024.01.25.577265

**Authors:** Benjamin Minch, Morgan Chakraborty, Sam Purkis, Mattie Rodrigue, Mohammad Moniruzzaman

## Abstract

Deep-sea brine pools represent rare, extreme environments that focus biodiversity at bathyal to abyssal depths. Despite their small size and distribution, brine pools represent important ecosystems to study because they provide unique insight into the limits of life on Earth, and by analogy, the plausibility of life beyond it. A distinguishing feature of many brine pools is the presence of thick benthic microbial mats which develop at the brine-seawater interface. While these bacterial and archaeal communities have received moderate attention, little is known about the viral communities and their interactions with host populations in these environments. To bridge this knowledge gap, we leveraged metagenomic and metatranscriptomic data from three distinct zones within the NEOM brine pool system (Gulf of Aqaba) to gain insights into the active viral ecology around the pools. Here, we report a remarkable diversity and activity of viruses of all nucleic acid types and genome sizes that infect prokaryotic and eukaryotic hosts in this environment. These include giant viruses (*phylum:* Nucleocytoviricota), RNA viruses, jumbo phages, and polinton-like viruses (PLVs). Many of these appeared to form distinct clades showing the possibility of untapped viral diversity in the brine pool ecosystem. Zone-specific differences in viral community composition and infection strategy were also observed with lysogenic phages seeming to dominate the bacterial mat further away from the pool’s center. Through host matching, viruses infecting metabolically important bacteria and archaea were observed – including a linkage between a jumbo phage and a key manganese-oxidizing and arsenic-metabolizing bacterium. Our findings shed light on the role of viruses in modulating the brine pool microbial community dynamics and biogeochemistry through revealing novel viral diversity, host-virus associations, and spatial-scale heterogeneity in viral dynamics in these extreme environments. These results will provide crucial foundation for further investigation into the adaptations of viruses and their microbial hosts in extreme habitats in the marine ecosystem.

## Introduction

Deep-sea brine pools represent one of Earth’s most extreme environments due to their hypersaline anoxic conditions, and low pH (1). Their formation is largely caused by the stable accumulation of hypersaline brine generated from the interaction between seawater and buried salt deposits which accumulate in seabed depressions (2, 3). So far, these unique environments are known to exist in only three major water bodies: the Gulf of Mexico, the Mediterranean, and the Red Sea (4–6). Within their respective bodies of water, brine pools are rare and small in comparison to their host basin, ranging in size from hundreds of square meters to a few square kilometers (7). Most Brine pools are found in deep-sea depressions, but a few have been found in shallower coastal-shelf waters of the Red Sea (8).

Though small, brine pools represent oases of life for both microorganisms and macrofauna. These environments stand in contrast to the rest of the deep benthos where life is scarcer than the photic zone (9,10). Brine pools teem with life, hosting dense microbial mats, as well as various species of bivalves, shrimp, and fish (7,11). Hence, brine pools have become key model ecosystems for studying the limits of life on our planet as well as the potential for life beyond it (12–14). Specifically of interest is the comparability of these deep-sea brine pools to the conditions of the subsurface oceans on the icy moons that orbit Jupiter and Saturn (12,15,16).

Early work advocated that these deep-sea brine pools were sterile, but this notion was drastically reversed after groundbreaking phylogenetic studies of the bacteria that thrive in association with the brine (17). The majority of microbial life in the brine pools exists at the interface between the anoxic brine and the surrounding seawater, as dense bacterial mats are often observed surrounding the pools (10,18–20). A hypothesized factor contributing to the abundant growth is the density gradient created at the brine-seawater interface that can act as a trap for organic and inorganic materials from seawater (21,22). Such trapping makes the brine a rich source of nutrients for specific microbes to harness for growth. Along with having a nutrient sink, the microbes that inhabit the sediment in and around a brine pool also exhibit vertical and spatial stratification due to different physiochemical gradients of chemicals such as manganese, sulfate, and potassium, promoting diverse microbial metabolic strategies (23,24). Another key aspect of the stratification of the microbial communities is the abrupt shift from oxic to anoxic conditions in the pool (25). This interface creates niches for a plethora of methanogenic and sulfate-reducing bacteria to flourish (26).

Bacterial communities in diverse environments are usually associated with the presence of viral populations, and it seems this holds true for even the most extreme of environments. Studies of brine pool sediments in the Mediterranean, for instance, demonstrate high levels of viral infection, proposing viral lysis as a key top-down control of prokaryotes in the brine pool sediments (27). This is not to say that eukaryotic grazing is not present, as zooplankton and fish have been observed feeding on the thick particles at the brine-seawater interface (7,28) and many species of fungi, dinoflagellates, and ciliates have been discovered around the pools (29).

Although viruses are typically thought of predominantly as cell-lysis agents, not all viruses adopt the same infection or replication strategy. Viral reproduction occurs primarily through either lytic or lysogenic infection, with the quantitative importance of each of these processes varying greatly throughout the ocean (30). Within the lytic cycle, viruses infect hosts, replicate inside of their cells and eventually lyse the cell to release more viruses into the environment. The lysogenic cycle involves a temporal separation between infection and lysis as the virus will integrate into the host genome. Viruses can switch between the lysogenic cycle and lytic cycles, a phenomenon called induction, which usually occurs from a variety of factors including changes in nutrients (31) or environmental stressors such as UV radiation (32). Bacterial populations associated with lysogenic viruses have been hypothesized to have a competitive advantage as lysogenic viruses may protect against infection from homologous viruses (33) and confer beneficial traits in the form of auxiliary metabolic genes (AMGs) (34). Although lysogeny was initially thought of as a survival strategy for viral communities with low host abundance (35), the recent “Piggyback-the-Winner” theory suggests that lysogeny predominates at high microbial abundance and growth rates due to the benefits of preventing niche invasion from other viruses (36).

Very few studies have delved into the viral communities in and around brine pools and none have focused on viral life strategies. In addition to the pioneering study of viruses in brine pools by Corinaldesi et al. (37), metagenomic-based viral community characterization has only been carried out in two studies using shallow sequencing approaches such as 454 pyrosequencing methods, all of which provide a preliminary analysis of viral diversity (38,39), focusing mainly on prokaryotic viruses. Besides the characterization of the majority of viruses belonging to the Caudoviricetes family (38), not much is known about the viral community inhabiting the brine pools. Since viruses are potentially the main regulators of microbial populations around the brine pools, this knowledge gap is an important one to fill to gain insight into the limits of life and viral complexity, as well as the possibility of selfish genetic entities like viruses influencing life elsewhere in the universe. A key knowledge gap is the lack of a clear assessment of the diversity, host interactions, and life strategies of viruses in brine pools, which is needed for evaluating their roles in structuring the microbial communities and biogeochemical cycles. Recent advances in viral metagenomics through high-throughput sequencing and methodological advances in the assembly and identification of viruses open the door to fill this knowledge gap to obtain a comprehensive understanding of the role of viruses in these extreme environments. In this study, we sought to investigate active viral ecology, leveraging both abundance (metagenomic) and activity (metatranscriptomic) data to inform viral life strategies and patterns of infection across three distinct zones within the most recently discovered Red Sea brine pools, the NEOM pool system (7,40). By integrating these two types of data and using newly developed viral identification and host linkage tools, we seek to elucidate the complexities of viral-host interactions across a horizontal spatial gradient around and within the NEOM brine pools. In addition to this goal, we also categorize the eukaryotic virus community, showing active eukaryotic viral infections at a water depth of 1,770m in an extreme hypersaline environment. Leveraging all of this information, we also investigate patterns in viral activity across the study zones to shed light on how viral ecological dynamics might conform to or deviate from those observed in marine environments.

## Results

### Sampling zones

The NEOM brine pool is the northernmost brine pool in the Red Sea (Figure 1a) and possesses three visually distinct zones within which surficial sediment samples were collected. Using the manipulator arm from a remotely operated vehicle (ROV) These zones include a “gray zone” (GR) and “orange zone” (OR) which are shoreward of the pool, as well as a “pool zone” (PL) which was sampled from the sediment inside of the brine pool (Figure 1b). Sediments from each of these zones were collected and DNA and RNA were extracted directly from the preserved sediments (See methods).

**Figure 1.**
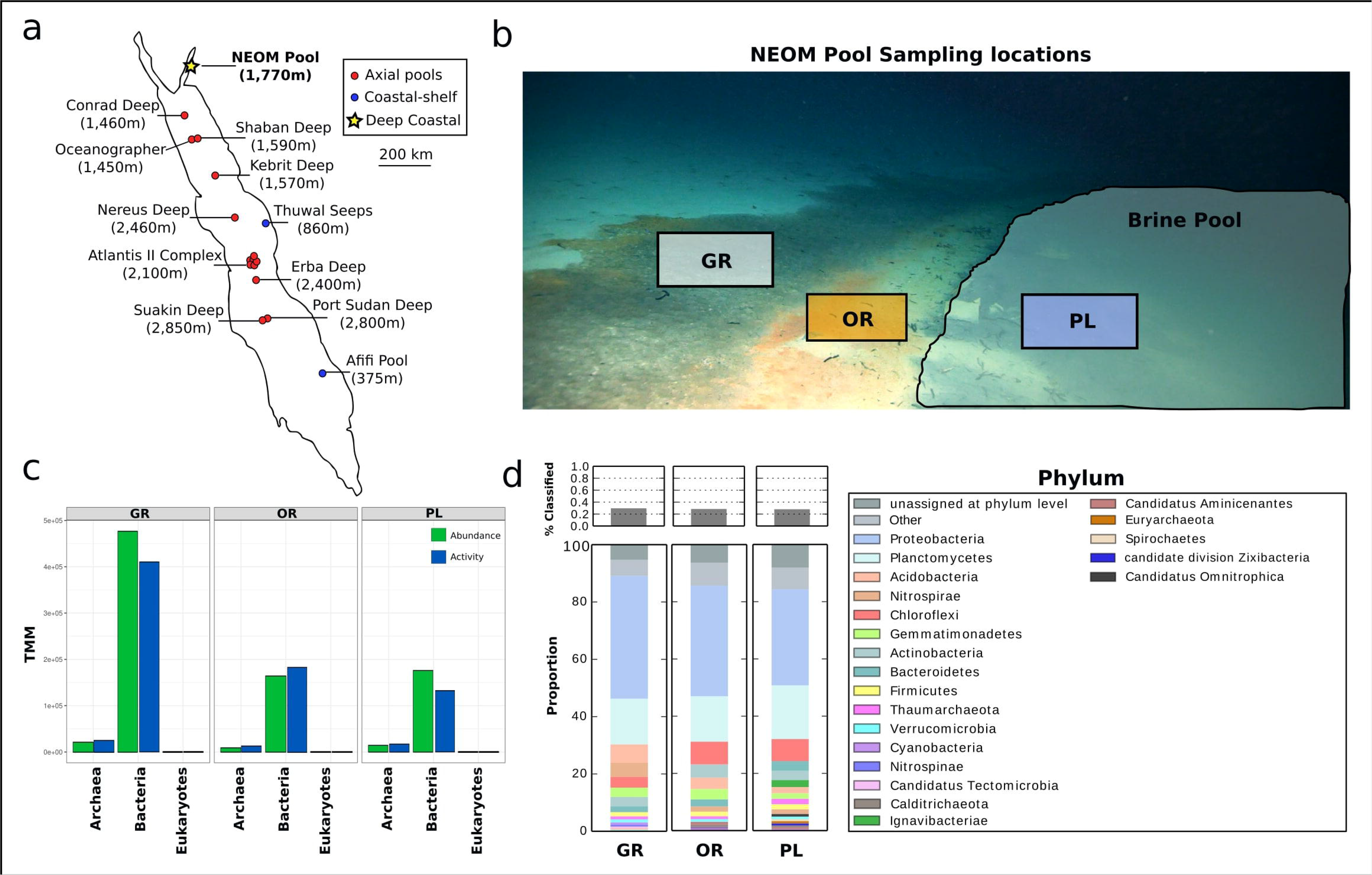
Sampling location and community composition. (**a**) A map of all discovered Red Sea brine pools, split into three categories based on location and depth. **(b)** A picture taken from the remotely operated vehicle (ROV) of the sampling locations showing the gray microbial mat (GR), orange mat (OR), and pool zones (PL). **(c)** Abundance and activity of different kingdoms of life based on normalized metagenomic (abundance) and metatranscriptomic (activity) reads (TMM) mapped to contigs from each kingdom identified using Tiara. **(d)** Relative abundance of different bacterial and archaeal phyla at each zone. These proportions were determined using kaiju to assign taxonomy to trimmed metagenomic reads.

### Prokaryote Abundance, activity and diversity

We estimated the abundance of prokaryotic communities in the study zones by mapping reads to the bacterial, archaeal, and eukaryotic contigs. This revealed an approximate 3-fold higher abundance of bacteria at the gray zone (GR) compared to the orange (OR) and pool (PL) zones (Figure 1c). Bacteria were the most abundant taxa present at all zones, making up between 92 and 96% of the total reads mapped at each zone. Archaea made up around 4-8% of total mapped reads at each zone and eukaryotes typically only made up ∼0.03% (Figure 1c). Of the 30% of bacterial reads that could be classified at each zone, the majority came from Proteobacteria (∼40%) and Planctomycetes (∼15%) across all zones. The gray zone had a higher abundance of Acidobacteria and Nitrospirae compared to the other zones, while the other zones had a higher proportion of Chloroflexi. The pool zone had a few unique taxa such as Ignavibacteriae and Euryarchaeota.

### Abundance, activity, and diversity of prokaryotic viruses

We used the ViWrap pipeline to identify a total of 1,184 viral genomes with various levels of completeness from the three samples. Genome sizes ranged from 1.3-427 kbp. To determine zone-specificity and spatial niche partitioning of the viruses in the brine pool, we established a cutoff criterion. In order for a virus to be deemed “exclusive” to a zone, it had to have a TMM-normalized abundance or expression value greater than 1 and over 10× higher values than the combined abundance or expression values from the other two zones. According to the established criteria, the gray zone had the largest proportion of unique viruses with 319 and 316 zone-specific viruses in the metagenomic and metatranscriptomic data (Figure 2a), respectively. More viruses were shared between the orange and pool zones than between either zone and the gray zone. There were also many viruses that were not abundant or expressed (TMM <1 for all zones), showing the presence of a rare virosphere. Overall, abundance (metagenomic) and activity (metatranscriptomic) followed each other closely, demonstrated by the high correlation between viral activity and abundance (Figure S1). Read mapping demonstrated a significantly higher diversity of viruses at the orange zone (p <2.2e-16) (Figure 2b) and a larger number of viruses present at the gray zone than the other zones (about a 4-fold increase), following the trend in prokaryotic abundances (Figure 2c). Interestingly, inside the pool was the only zone where activity (RNA) was comparatively higher than abundance (DNA), suggesting the presence of a highly active viral community inside the brine pool.

**Figure 2.**
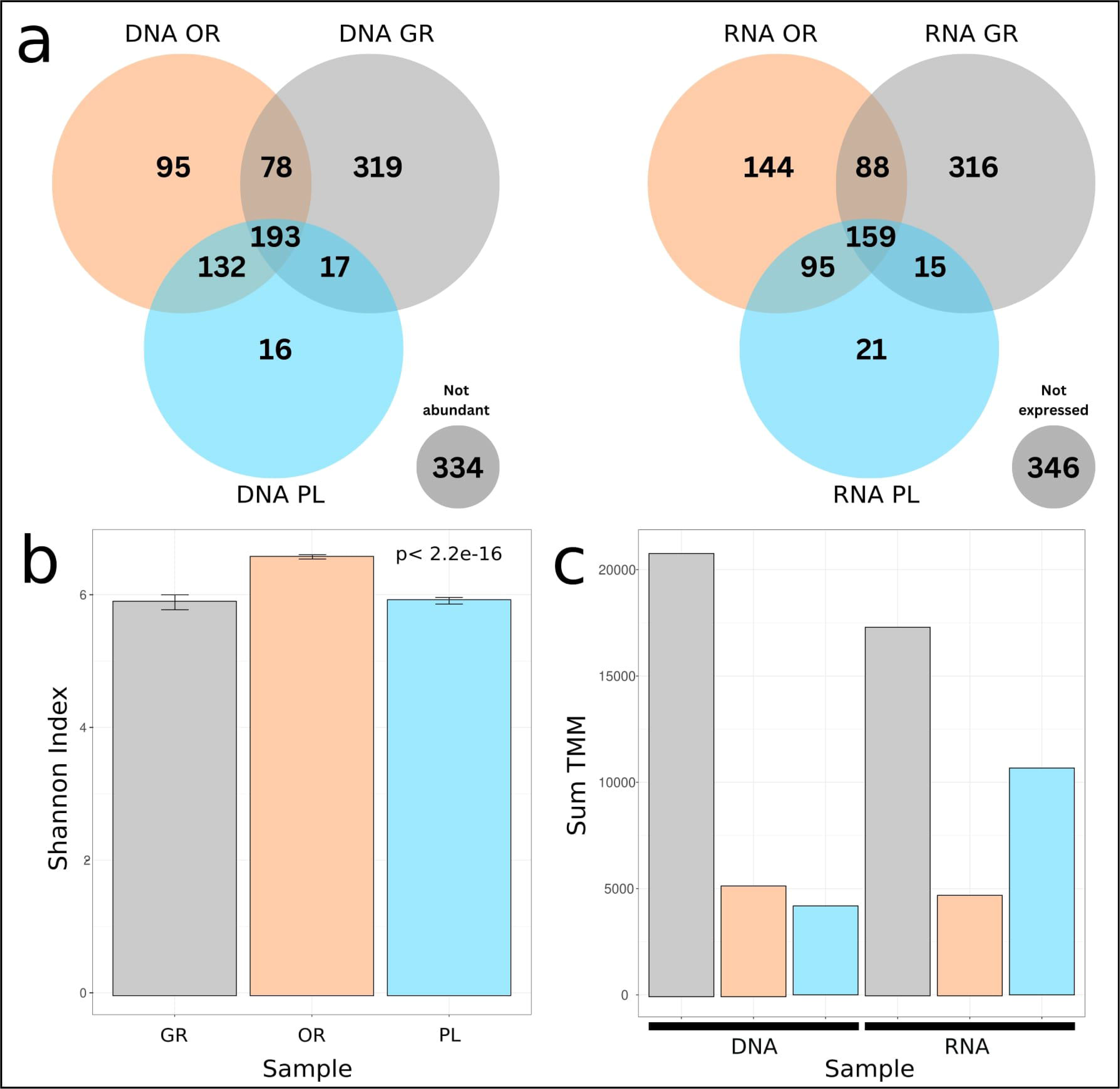
Zone specific patterns of viral community composition, diversity, and abundance. (**a**) Venn diagrams displaying unique viruses obtained within each zone for both metagenomic (DNA) and metatranscriptomic (RNA) datasets. A virus was determined to be specific to a zone if its abundance or activity was > 10 times greater than the other zones combined. “Not abundant” and “not expressed” viruses were determined by TMM values less than 1. **(b)** Shannon diversity of viral populations across zones. Error bars represent 2 SEM from the mean calculated from bootstrapping the population. P-value represents a t-test on the difference in mean between OR, GR, and PL zones. **(c)** Abundance and activity of viruses across zones (colored by zone). Values represent sum TMM of all viruses present at the zone.

Out of the 1,184 viral genomes, we were able to assign taxonomic labels to 258 (22%). These viruses made up around 25% of total viral abundance and activity at each zone, demonstrating the presence of a large viral dark matter. Of the viruses with taxonomic assignment, differences between zones emerged as the orange zone had the highest number of less abundant species, reflected in the higher diversity index. The orange zone had a higher proportional abundance of viruses showing homology to diverse known viruses including Bordetella phage, Rhodobacter phage, and Pseudomonas phage. The gray zone had higher proportions of viruses showing homology to Caulobacter phage CcrPW, Clostridium phage, Synechococcus phage Bellamy, and Xanthomonas phage. The pool zone had higher proportions of viruses showing homology to Croceibacter phage, Microviridae group, Paracoccus phage, Ruegeria phage, and Salmonella phage. All zones shared an abundance of viruses showing homology to Pelagibacter phage and Vibrio phage phiVC8 (Figure 3).

**Figure 3.**
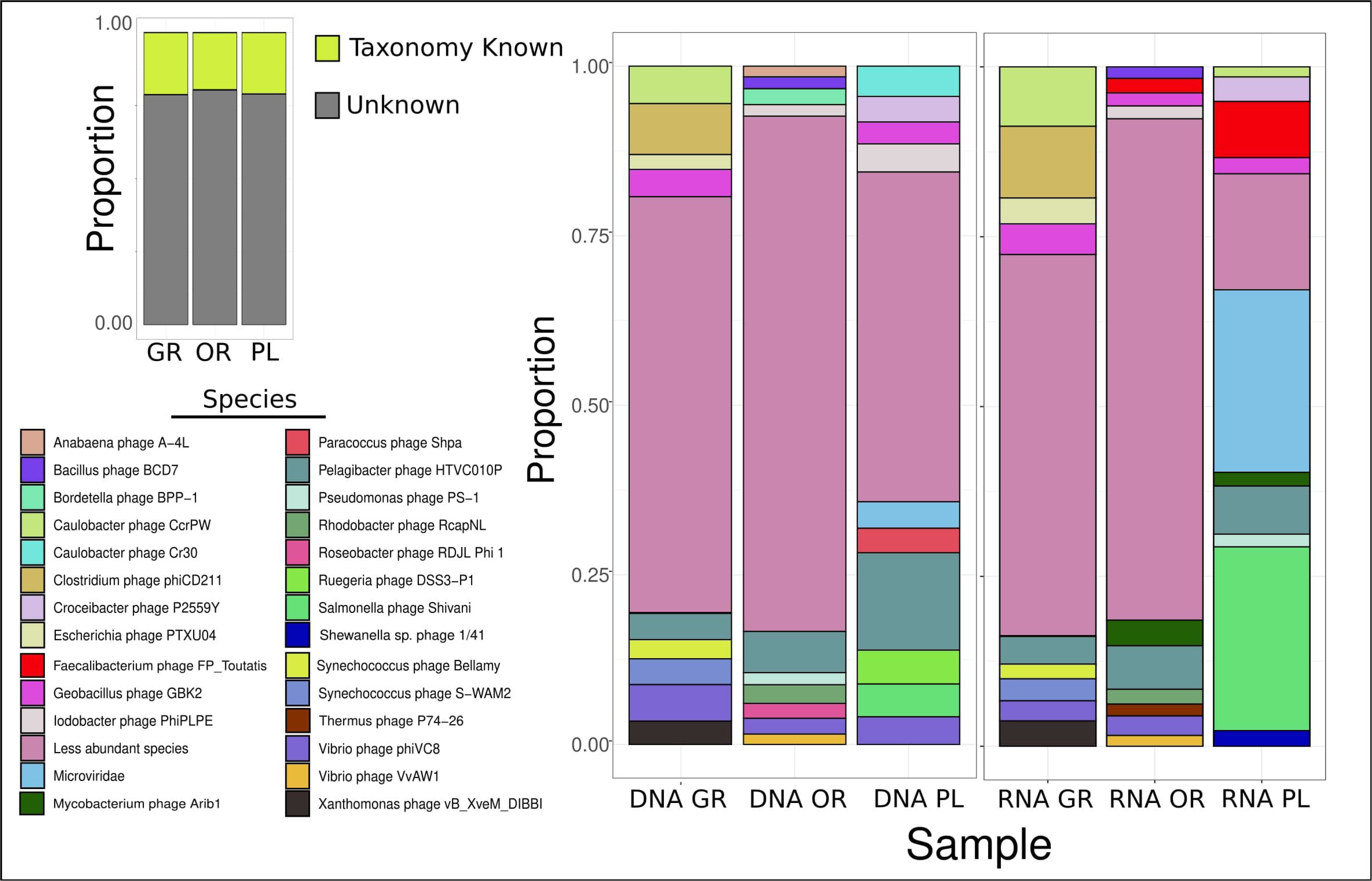
Viral community composition across zones. Viral homology at the species level was determined for 258 viruses, making up only ∼25% of the total abundance and activity of all viruses in the community. The proportion of the top 10 viral species for each zone is shown for metagenomic (DNA) and metatranscriptomic (RNA) data, with less abundant species being grouped together. Proportional abundance within a zone is determined from RPKM normalization, taking genome length and reads mapped as normalization factors.

The activity of viruses at both the gray and orange zones roughly matched the abundance data, but this did not hold true for the activity of viruses inside the brine pool. There is an overrepresentation of viruses with homology to Faecalibacterium phage FP_Toutatis (∼10% of total activity), Microviridae (∼25%), and Salmonella phage Shivani (∼25%). The high activity of these viruses in comparison with their abundance, combined with the higher total activity relative to abundance inside the pool, makes them likely to be the main contributors to this phenomenon.

### Eukaryotic virus abundance, activity and diversity

Viruses from the phylum *Nucleocytoviricota* (NCLDV), also known as giant viruses, are abundant in the Earth’s oceans and have been found to inhabit a range of different environments as well as modulate host metabolism and genome evolution (41,42). The main hosts for these viruses include microeukaryotes such as unicellular algae and protists. Searching for major capsid proteins (MCP) deriving from viruses from the phylum *Nucleocytovirocota* yielded a total of 33 MCPs in our brine pool data. Most of these MCPs belong to viruses in the *Imitervirales* order, but members of *Pandoravirales* and *Pimascovirales* were also identified (Figure 4a).

**Figure 4.**
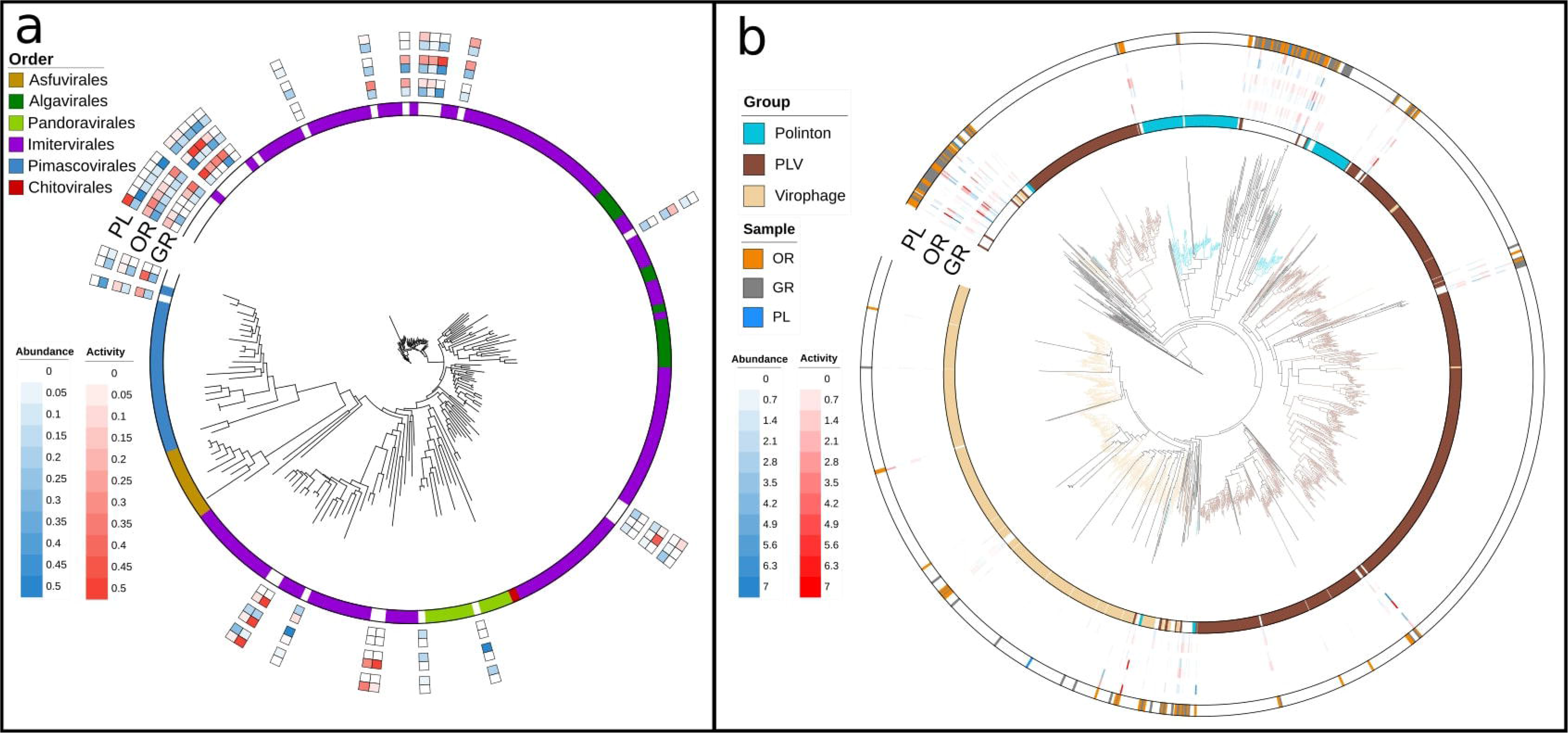
Phylogenetic diversity of eukaryotic viruses. (**a**) A phylogeny of the giant virus (NCLDV) major capsid protein (MCP) with reference sequences from the GVDB. Reads were mapped to MCPs and abundance and activity are represented for each zone as log normalized TMM values. (b) A phylogenetic tree showing identified MCPs from Polintons, PLVs, and Virophages from all zones as well as their abundance. Reference sequences were obtained from Stephens et al. (66). Reads were similarly mapped to MCPs and abundance and activity of individuals is shown in the heatmap.

NCLDVs were present across zones, with the orange zone having the largest total abundance (sum TMM-normalized abundance of 6.5) (Figure S2a). The gray zone had the lowest total abundance of NCLDVs (sum TMM-normalized abundance of 3.5). Most NCLDVs identified were found to be active, with the highest level of activity coming from *Imitervirales* in the orange zone. Although the one identified member of *Pandoravirales* was present at both the orange and pool zones, no activity could be detected at any zone. Other groups showed opposite dynamics, as members of the family Allomimiviridae (a family within *Imitervirales*), only showed activity data without appearing in the metagenomic data (Figure S2b). We also obtained a putative Algavirales member based on MCP phylogenetic analysis.

Virophages, Polintons, and Polinton-like viruses (PLVs) are viruses or virus-like transposable elements that have a shared evolutionary origin and can integrate into eukaryotic genomes or co-infect with NCLDVs. Recent studies have revealed that these viruses are likely common in diverse aquatic environments – however, their ecological roles and distributions in diverse extreme environments remain largely uncharacterized. These viruses all have distinct, yet evolutionarily related hallmark genes that can be leveraged for the discovery of sequences originating from these viruses. We used a set of major capsid protein (MCP) hallmark gene hidden Markov profiles we previously generated to search the brine pool sampling zones. This search revealed 183 viruses from these groups, consisting of Virophages, Polintons, and PLVs, indicating that these viruses are a key component of the eukaryotic virosphere in the brine pools. Phylogenetic analysis revealed that most brine pool sequences formed a new clade within the Polintons, with some also forming a deeply rooted group within the PLVs. (Figure 4b) Members of these putative new clades were highly abundant and active across all zones, with the highest activity at the orange and gray zones. Overall, identified sequences clustering with virophages were the least abundant and active of the three groups, but there was one identified MCP within this group that had activity at the orange and pool zones greater than all other elements at those zones combined (Figure S3).

We also identified numerous contigs representing RNA viruses from the metatranscriptomic data across all three zones (Figure S4). While representative RNA-dependent RNA polymerase (RdRp) sequences were found from all 8 major RNA virus phyla, the majority of sequences from the brine pool clustered into a putative novel clade. This clade was situated within the phylum Pisuviricota and the closest reference sequences belong to the Beihai picorna-like virus, with a known bivalve host, and the Ustilaginoidea virens non-segmented virus 2, with a known fungal host.

### Prokaryotic and putative Eukaryotic viral hosts

Despite the large diversity of novel viruses in these zones, we were able to predict hosts for 36 of the viruses (about 3% of the viral community) potentially infecting prokaryotes. The majority of these predicted hosts were members of the class Gammaproteobacteria, consistent with Proteobacteria being the most abundant phylum across zones (Figure 5a). A large proportion of host predictions were also from the Bacteroidia class. Zone-specific host associations were also observed such as Methanosarcinia and Nitrospinia being exclusively predicted at the gray zone, and Bathyarchaeia and Bacilli being exclusively predicted at the orange zone.

**Figure 5.**
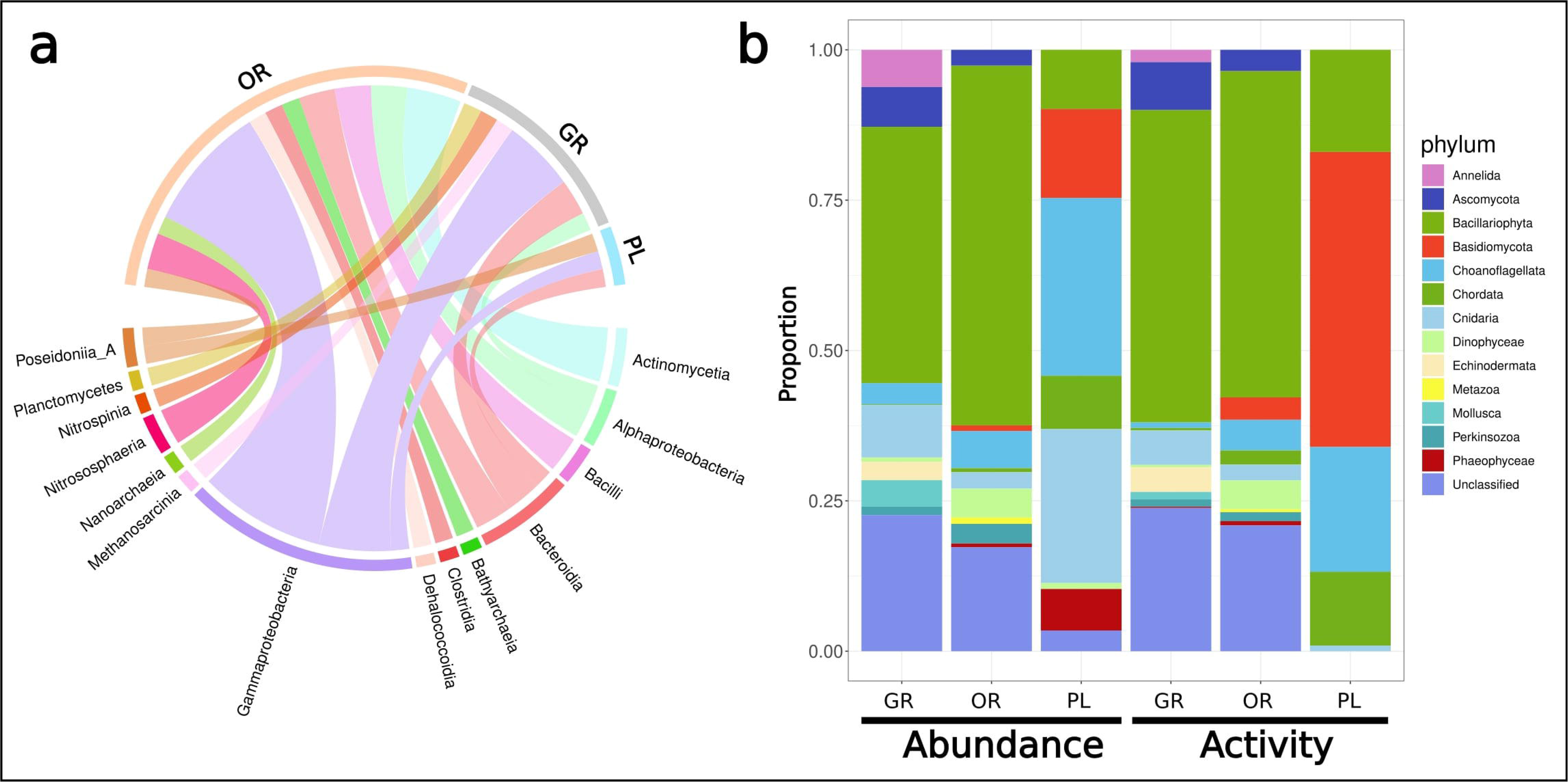
Predicted and potential hosts for prokaryotic and eukaryotic viruses. (**a**) Virus-host predictions for prokaryotic viruses were done using iPHOP for all zones. A total of 36 host predictions were made across zones and band thickness represents more hosts from a given bacterial class or zone. **(b)** The potential eukaryotic host pool was created through mapping reads to confirmed eukaryotic contigs obtained from Tiara. These contigs were classified to the phylum level using CAT and proportions represent RPKM values for each zone.

We also identified the eukaryotic community in the brine pool zones that potentially serve as the host pool of the NCLDVs present. Among the eukaryotic taxa present, the most abundant and active phylum around the edges of the pool was Bacillariophyta, making up over 50% of the abundance and activity at the orange zone and having similar high abundance at the gray zone (Figure 5b). Ascomycota, Choanoflagellata, and Cnidaria were all found to have relatively high activity and abundance at both gray and orange zones as well. Inside the pool, the dominant eukaryotic phyla were Basidiomycota, Choanoflagellata, and Cnidaria, although only the first two were represented in the activity data. Unique to the gray zone, Mollusca were also abundant and active on the outskirts of the pool.

### Active infection ecology of Brine Pool viruses

To further delve into the dynamics and functional potential of viruses in the brine pool, we separated the viral populations into four categories based on hierarchical clustering of individual virion rank activity to rank abundance ratios (Figure S5). A virus was deemed “non-abundant, non-active” if both activity and abundance were < 1, and these viruses were not included in the clustering method. Clustering revealed distinct groups of viruses based on their activity: abundance ratios in each of the zones. The gray zone had the highest number of “low-abundance, non-active” viruses (n=549) but also the highest number of “highly active” viruses (n=88) (Figure 6a). The orange zone had the highest number of “active + abundant” viruses (n=827), making up 90% of viruses in this zone. The pool zone was split nearly evenly between “active + abundant” and “abundant, non-active” viruses.

**Figure 6.**
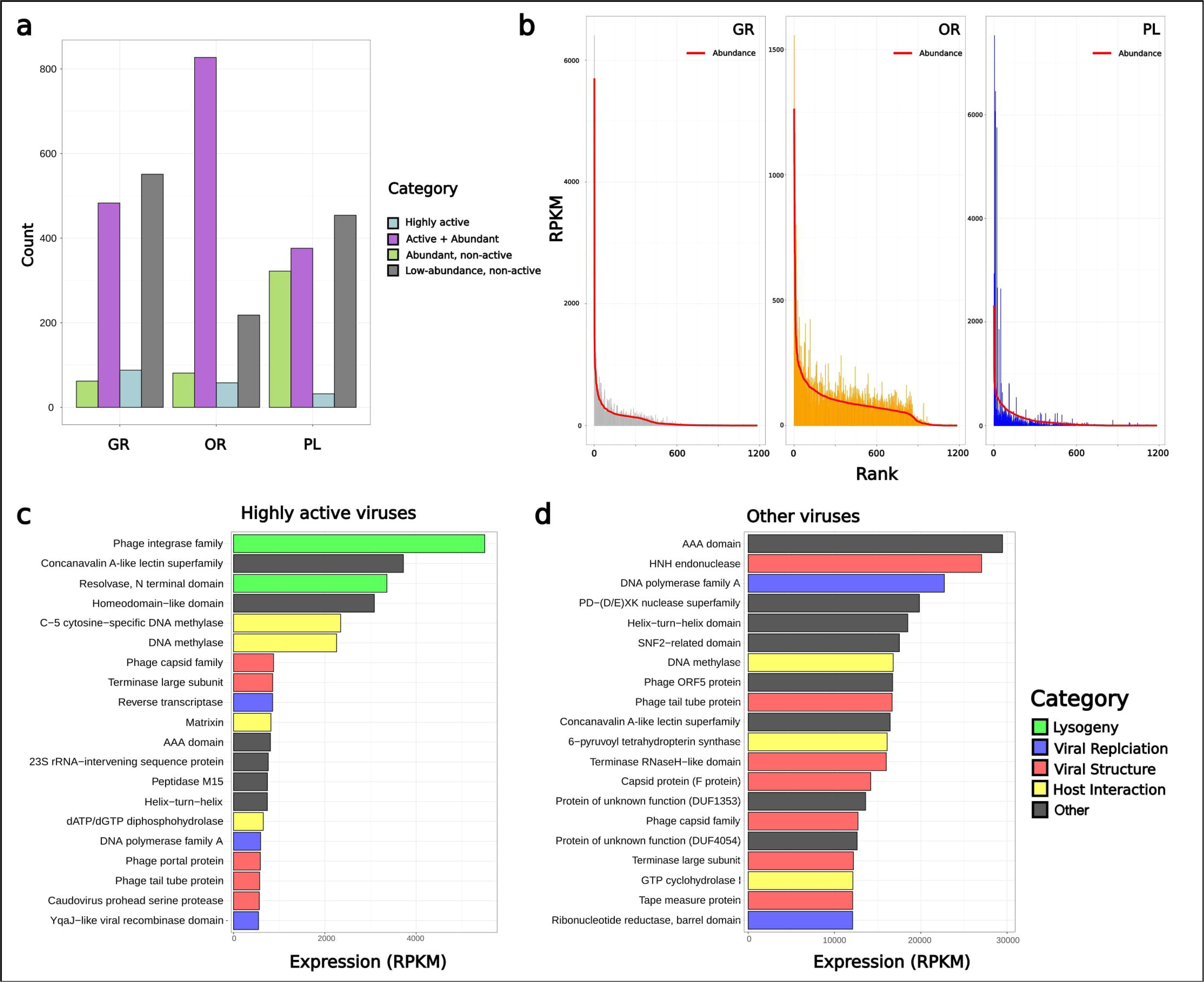
Active viral ecology and gene expression. (**a**) Viruses from each zone were categorized as either being “highly active”, “active + abundant”, “abundant, non-active”, and “non-abundant, non-active” based on hierarchical clustering of activity and abundance ratios. **(b)** Rank abundance curves for each zone with the red line representing metagenomic (abundance) read mapping and the colored bars representing metatranscriptomic (activity) data. Both read counts were normalized with RPKM. The gene expression profiles of **(c)** highly active and **(d)** other viruses. The top 20 genes (derived from sum RPKM) from each group are displayed in descending order. Categories were created based on gene annotations found within the PFAM database.

Analysis of the viral community’s rank abundance curves for both abundance and activity provides further insight into the differences in viral ecology across zones, confirming what was seen in Figure 6a and Figure S5 (Figure 6b). For all zones, there was a sharp drop-off in both activity and abundance with the rank abundance curve showing a strong left-skew, demonstrating that a few viruses at each zone made up a large portion of the activity and abundance for the given zone. However, this drop-off was less severe for the orange zone, and the rank abundance curve showed a more even distribution – with many viruses still showing high abundance and activity across ranks.

A total of 178 “highly active” viruses were identified across zones. In order to elucidate differences in gene expression patterns between these highly active viruses and other groups, the activity of the genes in these viruses was assessed through mapping of metatranscriptomic reads. Interestingly, while many genes crucial to viral replication and interaction with their host were similarly expressed across groups, the highly active viruses showed genes involved in lysogeny to be the most highly expressed (Figure 6c). This was not true of other viral categories, as genes related to viral replication and structure were among the most highly expressed (Figure 6d).

### Stratification of Gene Expression and AMGs across zones

To further understand differences in expression patterns between viral communities at the different zones, viruses specific to each zone were obtained (Figure 7a). These groupings were obtained from activity specificity analysis shown in Figure 2a. Once viruses were grouped into the three zone-specific groups, most expressed genes of that group were investigated to gain insight into the potential activity and life strategy of viruses at each zone (Figure 7b). The gray zone had everse transcriptase, integrase, and a type VI secretion system domain as its most expressed genes. Other notable genes of interest in this list are resolvase, a gene known to be involved in the lysogenic cycle, as well as a virulence-associated protein. An independent read mapping of phage integrase genes across zones (normalized for total viral activity and library sizes) confirmed this finding as the gray zone had a level of integrase expression nearly 3.5× higher than the orange zone, and over 13× higher than the pool zone (Figure S6).

**Figure 7.**
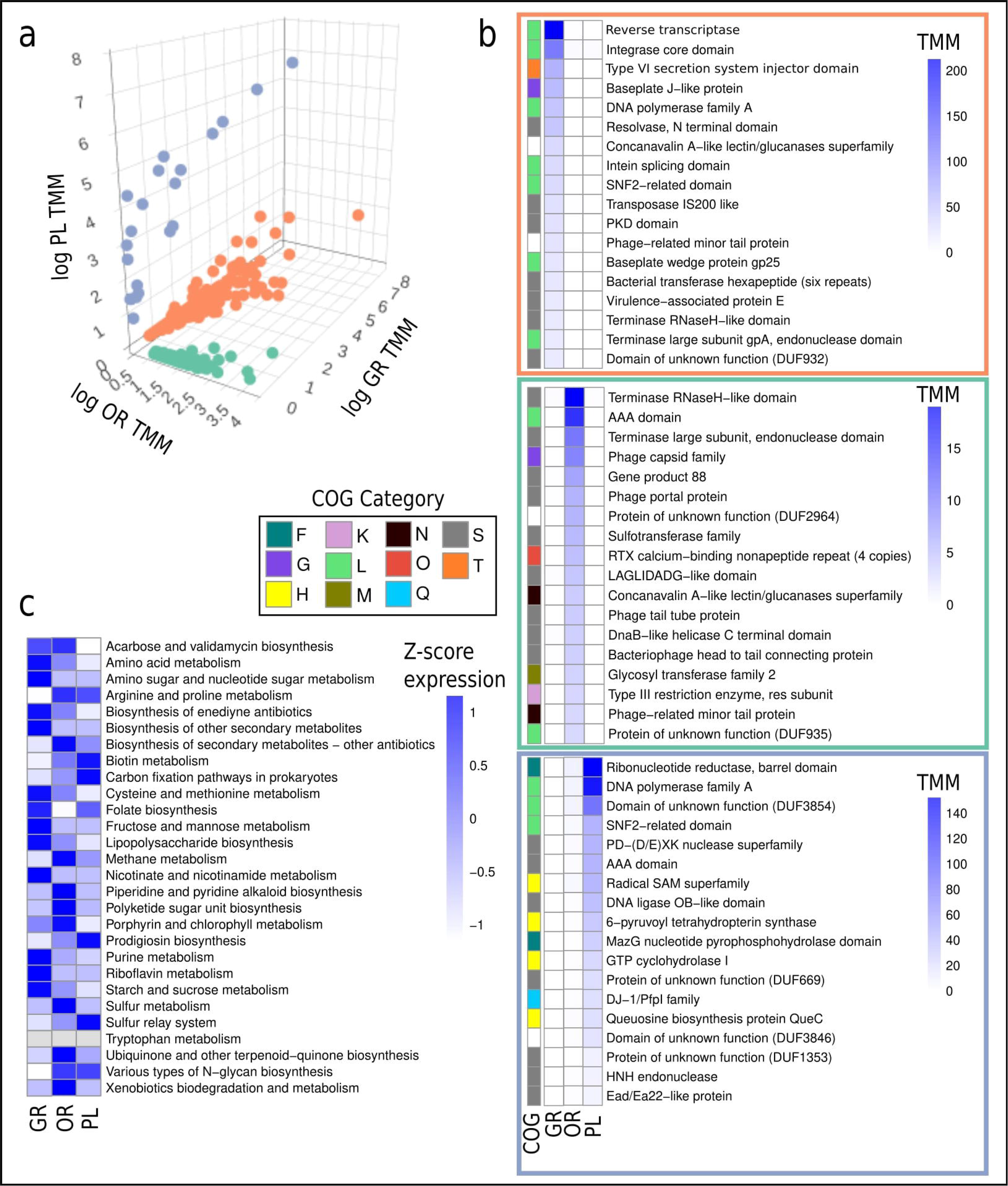
Zone specific marker genes and AMGs. (**a**) Viruses that were deemed “exclusive” to a certain zone (see figure 2) were displayed on a three-dimensional scatter plot with axes representing the log TMM of activity data from each zone. **(b)** Within each zone-specific cluster, the top 18 genes (by sum TMM) within that zone were plotted. COG categories were assigned based on eggNOG annotations [F: Nucleotide metabolism, G: Carbohydrate metabolism, H: Coenzyme metabolism, K: Transcription, L: Replication and repair, M: Cell wall/membrane biogenesis, N: Cell motility, O: Post-translational modification, Q: Secondary metabolite biosynthesis, S: Unknown, T: Signal transduction]. **(c)** Auxiliary metabolic genes (AMGs) were obtained from viruses at each zone and reads were mapped to calculate activity. This heatmap shows across-zone normalized Z-scores from TMM values.

The orange zone has terminase-related genes and an AAA domain-encoding gene (both genes involved in phage DNA packaging) as its most highly expressed along with the phage capsid. Other genes of interest that were highly expressed were sulfotransferase, the RTX calcium-binding repeat, and a glycosyl transferase family 2 gene. The pool zone had genes involved in replication among the most highly expressed ones such as ribonucleotide reductase, DNA polymerase A, and an SNF2-related protein. Among the most highly expressed genes at this zone, many were involved in coenzyme transport and metabolism including a gene belonging to the Radical SAM superfamily, 6-pyruvoyl tetrahydropterin synthase, GTP cyclohydrolase, and QueC, a gene involved in queuosine biosynthesis.

In total, 101 different auxiliary metabolic genes (AMGs) were recovered from the viral genomes, belonging to 28 different metabolic pathways (Figure 7c). Some of these AMGs recovered are homologous to genes from pathways involved in the biosynthesis of important biomedical compounds such as acarbose, validamycin, and enediyne antibiotics. Other interesting pathways include the recovery of genes involved in methane (mgsA) and sulfur (cysH) metabolism as well as xenobiotics degradation (guaA) which are most highly expressed in the orange zone. On average, pathways involved in amino acid metabolism and synthesis were more highly expressed in the gray zone (genes from 4 of the 5 pathways present).

### Jumbo phages infecting metabolically relevant hosts

Our genome binning approach implemented in the ViWrap pipeline recovered the genomes of two jumbo phages (phages with genome sizes larger than 200 kbp), and we were able to link both viruses to bacterial hosts through CRISPR spacer matching (Figure 8a). One jumbo phage with a genome size of 406 kbp was recovered from OR and one with a genome size of 204 kbp was recovered from GR. These two jumbo phages showed high abundance and activity, ranking within the top 1% in terms of activity and abundance at their respective zones (Figure S7a). Both jumbo phages also showed high zone-specificity, with activity and abundance levels at their zone of recovery over 11× and 184× higher for the OR and GR jumbo phages, respectively (Figure 8b). These zone-specific jumbo phages also contained unique genes, not present in the other jumbo phages, such as DNA methylase and dockerin encoded in the orange zone jumbo phage genome, and CHC2 zinc finger domain-containing protein, and cell wall hydrolase in the GR jumbo phage (Figure S7b).

**Figure 8.**
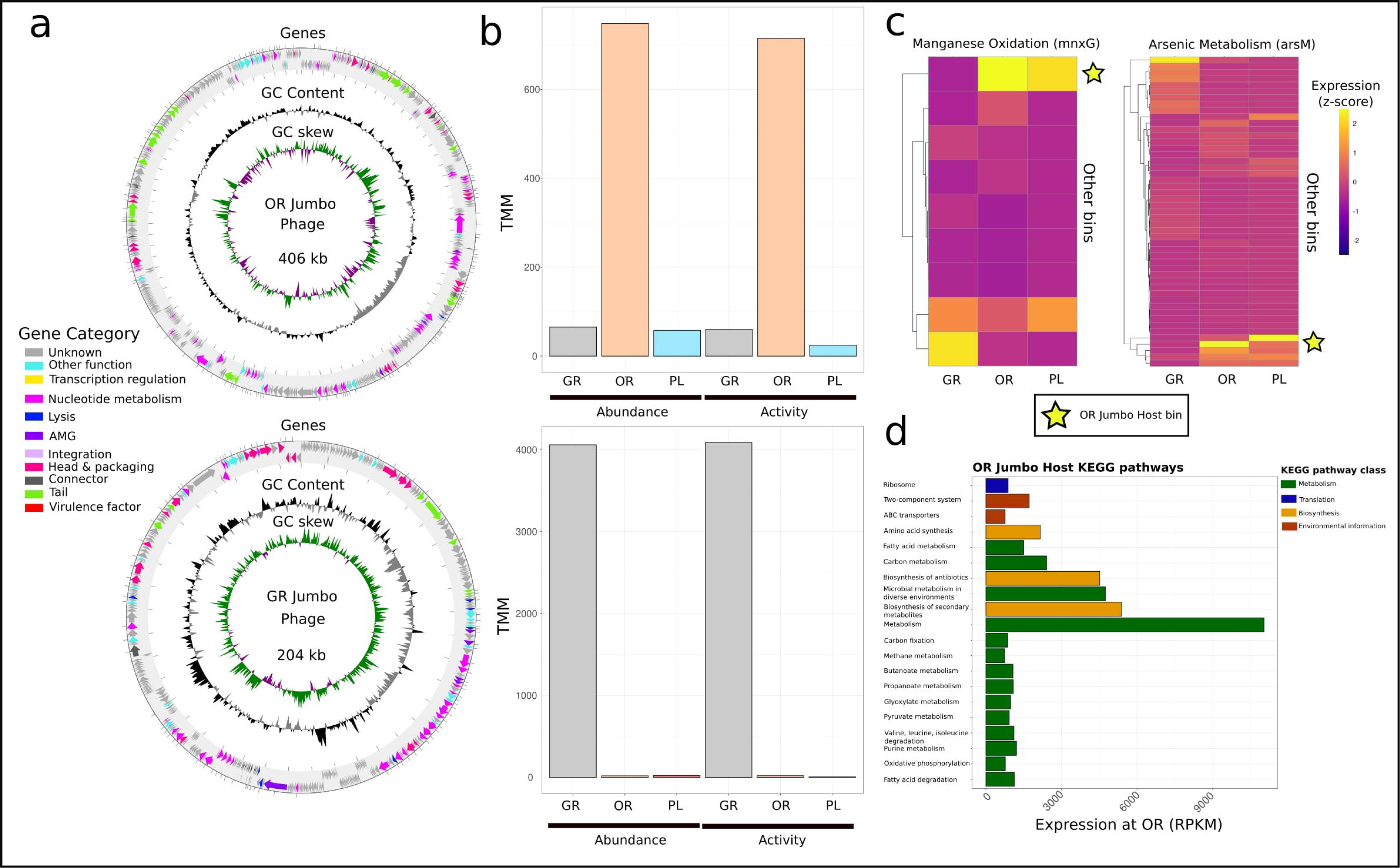
Jumbo phages and their hosts. (**a**) Genome maps of two identified jumbo phages were created using pharokka. GC skew, GC content, and gene categories were displayed on each track. **(b)** Reads were mapped to each jumbo phage genome and normalized using TMM across zones. Abundance and activity for each phage at each zone is represented here. The metabolic profile of the CRISPR-linked host of the OR jumbo phage is displayed through both a targeted and general approach. **(c)** Manganese oxidation and arsenic metabolism gene expression from all bacterial bins was calculated through read mapping to recovered genes and TMM normalization. Normalized reads were displayed as z-scores, normalized for within zone comparison to parse out key players in each process. The stars represent the OR jumbo host bacterial bin’s expression of these respective genes. **(d)** Reads were mapped to genes predicted from the OR host bacterial bin and normalized using RPKM. KEGG pathways were put into larger self-determined categories based on similar functions.

Through CRISPR spacer matching between these jumbo phages and bacterial bins recovered from the brine pool, we were able to match both jumbo phages with a potential host. While the potential host for the GR jumbo phage was a Proteobacteria with a partial genome, the host for the OR jumbo phage was found to be a metabolically active bacterium from the phylum *Myxococcota*. This bacterium was found to be the highest contributor among bacterial bins to manganese oxidation and arsenic metabolism at this zone, having the highest expression of genes involved in these pathways (Figure 8c), and contains genes involved in methanogenesis and biosynthesis of antibiotics that are expressed (Figure 8d).

## Discussion

### Spatial stratification of the bacteria and virus communities

Through leveraging a multi-omics approach, we were able to gain insight not only into the diversity and abundance of viruses and their potential hosts, but also into their active infection dynamics, gene expression, and putative life strategies. It was surprising that even across spatial distances of just a few meters, bacterial and viral communities were stratified, displaying large differences in both diversity and abundance. Previous studies have demonstrated vertical stratification of brine pool bacterial (23) communities, but here we show that within short distances in the same pool, the viral community varies dramatically. Our data is evidence that stratification is not only caused by the oxic/anoxic gradient, but also by an underlying nutrient or physiochemical gradient along the edge of the pool. Other studies have characterized potential gradients of methane, salinity, and temperature around brine pools which could be contributing to the stratification seen here (43–45).

Bacterial abundance and viral abundance followed similar patterns, with the gray zone having the highest abundance of both. A potential explanation for these patterns comes from the fact that while the brine is a nutrient-rich particle trap, the brine is also a hypersaline pool of water that restricts the growth of many microbial species. The brine has also been reported to splash outside the pool with minor disturbances and is episodically disturbed by underwater landslides (7, 40). The gray zone could be situated in the perfect area that can receive vital nutrients from the brine pools without too much direct contact with the harmful brine. This “sweet spot” for growth would be able to support the large bacterial abundance seen at the gray zone, nearly 3× more than other zones closer to the pool. In this way, the brine pool ecosystem can be thought of like the intertidal zone. Within the intertidal, the occasional waves of brine bring both nutrients and potential dangers of being inundated (46).

### The viral dark matter of the NEOM brine pools

Previous studies of viruses in the brine pools have found high levels of unclassified prokaryotic viruses within the pools (38,39) and our study confirmed this, with a large proportion of viruses showing no homology to known representatives. From the viruses that did show homology to known representatives, it is notable that only a small proportion of the viruses contribute to the bulk of activity at the pool (PL) zone. Viruses showing homology to Microviridae, and Salmonella phage Shivani were the main contributors to viral activity at this zone. Microviridae has been reported in deep sea environments before (47), but this is the first report of this group of viruses in a brine pool.

In addition to finding a large diversity of prokaryotic viruses, novel eukaryotic viruses were identified around the NEOM brine pool. Previously, viruses of the phylum *Nucleocytoviricota* have been found in metagenomic data from the brine pools (38), but the conclusion was that these viruses were most likely “pickled” (preserved in the hypersaline brine but not active) and not actively infecting hosts. Here, through leveraging metatranscriptomic data, we demonstrate that NCLDVs are not only abundant in the brine pools, but are also active, possibly infecting eukaryotic hosts. While it is true that some NCLDVs could be pickled due to lack of metatranscriptomic signal, the majority of recovered NCLDVs were both active and abundant.

While there is no established host-prediction tool for NCLDVs like phages, the host pool is quite limited due to the low eukaryotic diversity in the brine pools compared to productive surface waters (10). Among the potential hosts, members of the phylum Bacillariophyta (Diatoms) dominated the activity and abundance of the gray and orange zones. While there has never been a confirmed infection of Diatoms by NCLDVs *in situ,* many studies have noticed connections between these two groups through HGT networks and single-cell host approaches establishing cross-linkages (48–50). Other potential eukaryotic hosts for these viruses include Mollusks (51), Dinoflagellates (52), and Phaeophytes (brown algae) (53) – all of which have been previously observed in lab studies. Fungi could also serve as potential hosts and NCLDV-fungi host associations were previously reported as suggested by Bhattacharjee et al. (54) and Moniruzzaman et al. (55).

In addition to NCLDVs, we identified many Polintons, Polinton-like viruses (PLVs), and virophages in and around the brine pool. These evolutionarily related groups (56) all infect eukaryotic hosts and are widely abundant in aquatic ecosystems and within the genomes of eukaryotes (56,57). The presence and activity of virophages also confirm the activity of NCLDVs in the brine pools as virophages typically depend on NCLDVs for co-infection (58). The formation of a novel clade within the Polintons shows the potential untapped viral diversity within extreme environments. These results underline the importance of further expansion in our understanding of the role of Polintons, PLVs, and virophages in shaping eukaryotic evolution and the implications of their interactions in shaping the microbial community in diverse extreme environments the limits and potential origins of life in the deep sea (59).

Recent advances in cataloging the diversity and abundance of RNA viruses in the world’s oceans (60,61) suggest that the ocean is a large reservoir of untapped RNA virus diversity. These viruses are seen as important for exploring the origins of life under the RNA world hypothesis as they possess ancient genes such as RNA-dependent RNA polymerase (RdRp) (62,63). The finding of RNA viruses from 8 major phyla around the brine pool, as well as the discovery of a potential new clade of viruses within the Pisuviricota, indicates that extreme environments, like deep-sea brine pools can harbor novel RNA virus diversity, with our study representing the first identification of RNA viruses within brine pool environments. This active RNA virus community is likely supported by eukaryotic hosts such as fungi and mollusks; both of which were found in our metagenomic data. Future in-depth exploration of these viruses in brine pools and other extreme environments will help shed light on their adaptation in such environments and their possible ancient origins, assisting in evaluating the possible scenarios into the origin of early life.

### Active viral ecology at the brine pool

Besides characterizing the virosphere of the brine pools, we also sought to gain insight into virus-host dynamics and life strategies in an extreme environment. Through leveraging metagenomic and metatranscriptomic data, we were able to show that a small number of viruses at both the gray (GR) and pool (PL) zones were contributing to the majority of total viral activity and abundance at these zones. This is reflected in viral diversity as it was significantly lower in these two zones compared to the orange zone, which displayed a large number of viruses that were both active and abundant. These differences can be explained in part by the adherence to the viral ‘bank’ model which suggests that at a given time only a fraction of the viruses in a zone are active, while the majority are inactive (64). While the GR and PL zones seem to obey the model, the OR zone deviates, demonstrating the possibility for the breakdown of generalized viral strategic models when applied to specific individual environments. The pool zone also had a much larger number of viruses that were abundant but not active, demonstrating the possibility of the increased “pickling” effect inside the brine pool, where virus-host interactions might be influenced by the extreme salinity.

Our categorization of virus communities based on their level of activity helped to elucidate the infection strategies of the viruses that were highly active relative to their abundance. Analysis of gene expressions revealed that many viruses in this highly active category were most likely temperate phages. Further exploration into lysogenic markers such as integrase showed that the majority of these temperate phages are likely most active at the gray zone, furthest away from the brine. This zone has the highest bacterial population, which seems to be consistent with the “Piggyback-the-Winner” hypothesis that predicts temperate strategies become more prevalent with higher bacterial abundance (36).

In addition to elucidating viral infection strategy, the metatranscriptomics data also gave insight into the potential metabolic impacts these viruses may have on their hosts. Previous studies have found high methane concentrations in several brine pools (65–67). Here we find that viruses around the brine pool infect key methanogenic bacteria and archaea such as Methanosarcina (68) as well as contain AMGs within their genomes that aid in methane metabolism (msgA). These findings suggest that viruses play a key role in regulating and modulating methane metabolism in the brine pool environment as has been suggested in other environments (69,70). Genes for sulfur metabolism (cysH, mec), another key metabolic pathway in the anoxic brine pools (8,71), were also found among viruses from all zones.

In addition to providing added metabolic functionality, there is also evidence that viruses in the brine pool could be conferring helpful genes for stress tolerance. The high expression of genes such as GTP cyclohydrolase, QueC, and the Radical SAM superfamily gene at the pool zone suggests that these could be helping with anoxic and hypersaline stress (72–74).

Brine pools are unique environments that have been shown to be enriched in manganese and arsenic (7,75). This enrichment compared to the surrounding sediment allows for specific prokaryotic adaptations to utilize these trace elements (76). Through direct CRISPR spacer linkage, we identified a jumbo phage infecting one of these key microbial players in manganese oxidation and arsenic metabolism. This linkage presents the possibility that deep sea jumbo phages could be regulating key biogeochemical cycles important to life in the brine pools. The recovery of complete jumbo phage genomes from the brine pool adds to an expanding understanding of the biogeography and potential functions of marine jumbo phages (77).

### A conceptual model of brine pool virus ecology

Based on the patterns in viral diversity and ecological strategies observed in our study, we propose a hypothetical framework to explain the complexities of viral ecology within brine pools (Figure 9). This framework is grounded on the premise that brine pools behave similarly to intertidal habitats in that they show spatial stratification due to sediment physicochemical differences and the splashing of nutritious but potentially hazardous brine out of the pool (40). The combination of these two factors creates unique microhabitats, allowing for unique viral-host interactions and abundance patterns across a small scale of stratification.

**Figure 9.**
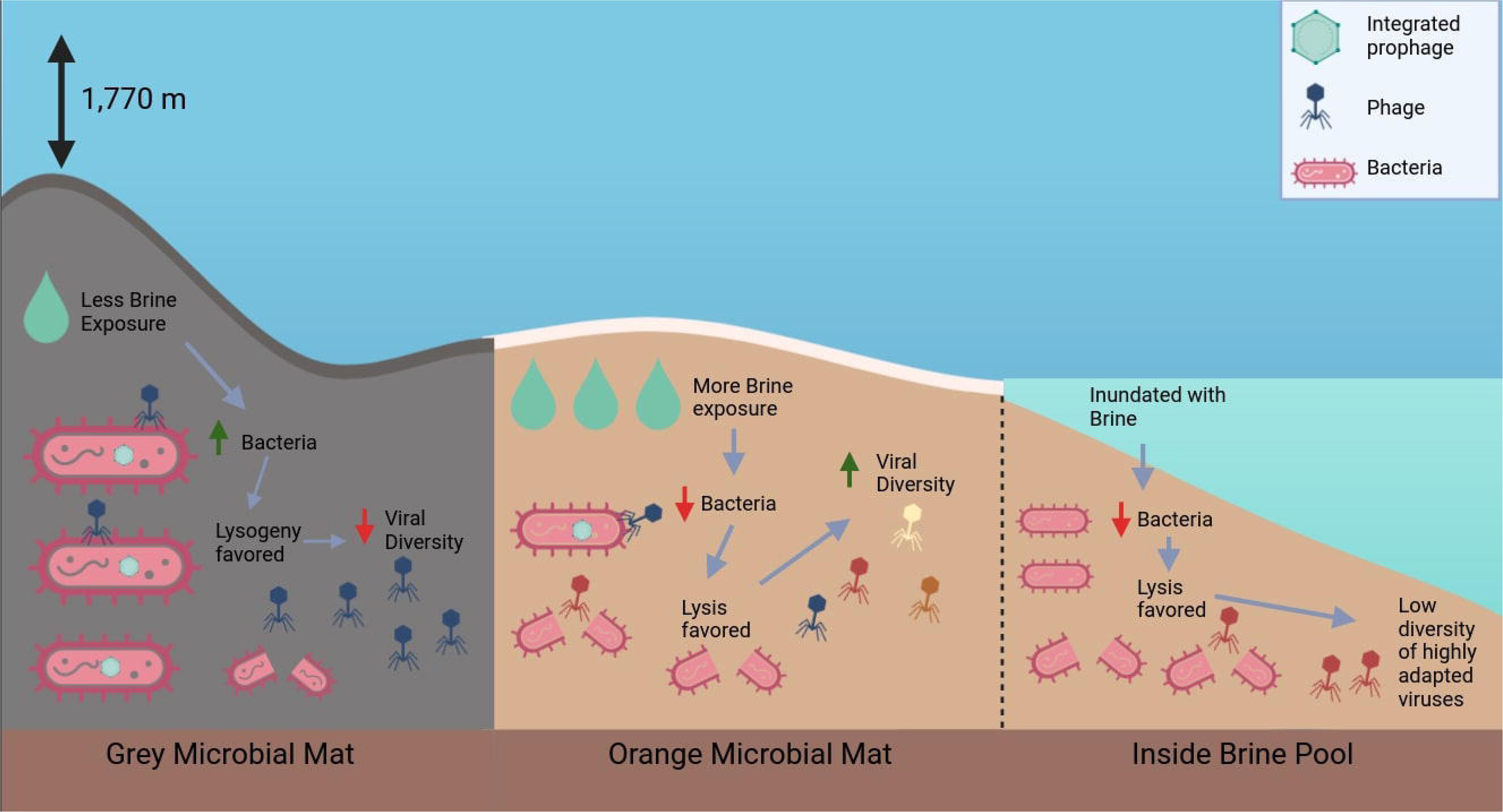
A conceptual model of brine pool virus ecology. An artistic rendition of our hypothesized prokaryotic viral ecology in the brine pool.

The gray zone, which is furthest away from the center of the pool but still close enough to benefit from nutrients delivered by it, shows the highest bacterial abundance. This high bacterial abundance leads to an increase in lysogeny among the viral population, as high bacterial density and abundance are key factors in predicting lysogeny in the “Piggyback the winner” model (36). The higher presence of lysogeny at this zone causes an increase in “superinfection exclusion”, a phenomenon in which lysogeny of one virus prevents others from infecting the same host (78,79). This exclusion may explain why in this zone, there is a lower diversity of viruses and only a small proportion of the viral community that contributes to overall viral activity and abundance. However, in the absence of additional data, it is important to note that factors other than superinfection might also contribute to the observed low diversity of viruses in this zone.

At the orange zone, right at the interface of the brine pool, microbes are likely exposed to the brine daily either through soil penetration or splashing (7,40). This increased exposure may be responsible for observed lower bacterial populations, favoring viruses undergoing the lytic cycle (80). With the lytic cycle favored, less superinfection exclusion takes place and there is room for more competition, explaining higher viral diversity at this zone. The increased competition and diversity of contributing viruses at the orange zone could also explain the observation of high expression of key metabolic genes such as those involved in sulfur and methane metabolism. The ability of a virion to confer a metabolic advantage to its host may confer a selective advantage, leading to an increased importance of AMGs in this zone (81).

Inside the brine pool, a zone characterized by full hypersaline and anoxic conditions, lower bacterial abundance is observed. Similar to the orange zone, lytic viruses are favored but unlike the orange zone, there is a lower viral diversity due to the potential selection of fewer highly-adapted viruses such as Microviridae. These highly-adapted viruses likely have a large impact on host growth and survival in this extreme zone (82).

We acknowledge that our hypothetical framework is based on the data from this single study, and there might be room for alternative explanations for the patterns we observed. However, this hypothetical framework provides the foundation for future in-depth research on the viral dynamics of diverse brine pool ecosystems to tease apart their role in biogeochemistry and niche differentiation of microbial communities.

## Conclusion

Overall, our study aimed to address the knowledge gap regarding the viral community in extreme brine pool environments. Through integrating metagenomic and metatranscriptomic data we were able to elucidate viral strategies and infection dynamics at a spatial scale in a brine pool Our data provides valuable insights into the viral community within these pools and sets the stage for further exploration and characterization of the physicochemical parameters that potentially drive the observed stratification in viral diversity and ecological strategies. Results from this research will be instrumental in generating further hypotheses and exploration into these unique environments with flourishing life around them.

## Methods

### Sample Recovery

The most recently discovered brine pool cluster in the Red Sea, the NEOM brine pools, was discovered in 2020 (7). These pools are distinct from previous Red Sea discoveries in that they were found only 2 km from the coast at a depth of 1,770m. This brine pool was found to have temperatures between 21 and 22 C, as well as a salinity of 160 PSU. (7). Another recent geochemical analysis of the NEOM brine pool found elevated levels of K^+^, Ca^2+^, Sr^2+^, B, Mg^2+^, and Li^+^ inside the pool compared to surrounding water (40).

During a 2022 research mission conducted and facilitated by OceanX, samples were collected aboard the R/V OceanXplorer in the Gulf of Aqaba, Red Sea. Using an Argus Mariner XL remotely operated vehicle (ROV), 3 scoops of about 500g of sediment were collected in each of the microbial zones surrounding the NEOM brine pool (gray, orange, and inside the pool) and sealed into separate quivers for ascent. Once secured aboard the research vessel, the sediment scoops were subsampled into 15 ml falcon tubes and preserved in RNALater. All samples were kept at 4°C until further analysis.

### Nucleic acid extraction and Sequencing

The DNA extractions were performed with the MagMax Microbiome kit from ThermoFisher following the manufacturer’s protocol. RNA was extracted with the RNA Powersoil Total RNA kit from Qiagen.

For sequencing, metagenomic libraries were prepared with the xGen ssDNA & low-input DNA prep kit from IDT. For metatranscriptomic libraries, ribosomal RNAs were removed with a 5S/16S/23S Fastselect kit (Qiagen). These libraries were prepared with the Kapa Hyper Stranded mRNA library kit (Roche). All samples were pooled and run on a NovaSeq 6000 (Illumina) with V1.5 sequencing kits for 151 cycles. Generated FASTQ files were demultiplexed with bcl2fastq (v2.2) conversion software (Illumina). The generated paired-end reads were 150 bp in length.

Metagenomic and metatranscriptomic raw reads were trimmed using TrimGalore, which uses Cutadapt and FastQC to trim and remove poor quality bases (v0.6.10)(83), and assembled using Megahit (v1.2.9) with the ‘-meta’ flag (84). Trimming and assembly statistics are present in a supplemental table (Table S1).

### Prokaryotic and Eukaryotic Abundance and Diversity

In order to assess bacterial and archaeal abundances, bacterial and archeal contigs >10kb were identified with Tiara (v1.0.3)(85) using a probability score cutoff of 0.65. Contigs were de-replicated at 95% average nucleotide identity (ANI) using dREP (v3.4.5)(86) resulting in a total of 266,243 bacterial contigs, 15,017 archaeal contigs, and 47 eukaryotic contigs. Trimmed reads were mapped to these contigs at 95% minimum identity using minimap2 (v2.17)(87) and coverage was determined by CoverM (v0.6.1)(88). Raw counts were normalized using the trimmed mean of M-values (TMM) method in edgeR (v3.18)(89) to take into account differences in library size and contig length. For each zone, these values were summed for all bacterial, archaeal, and eukaryotic contigs and plotted using ggplot (v3.4.4)(90). Metagenomic data was used to determine abundance and metatranscriptomic data was used to assess activity.

Bacterial diversity was determined using kaiju (v1.9.2)(91) with trimmed metagenomic reads from each zone. Taxonomy for each read was determined using the NCBI nr database (92) with a low abundance filter of 0.5 and a subsample percentage of 10. Eukaryotic contigs were classified to the phylum level using CAT (v5.3)(93).

### Prokaryotic virus taxonomy and diversity

Prokaryotic viruses were recovered from metagenomic datasets using the ViWrap (v1.2.1) pipeline (94), which uses both VIBRANT (v1.2.1)(95) and VirSorter (96) to identify viral contigs, and vRhyme (v1.1)(97) to bin viral contigs into metagenome-assembled genomes (MAGs). Viral MAGs were classified as part of the pipeline using vContact (v0.11.3)(98) to find viruses of the nearest homology. After running the ViWrap pipeline, resulting viral MAGs were dereplicated at 95% average nucleotide identity (ANI) using dRep. This resulted in a total of 1,184 viral genomes.

Trimmed metagenomic and metatranscriptomic reads were mapped to viral genomes at 95% minimum identity using minimap2 and coverage values were generated using CoverM. Raw read counts were normalized using TMM normalization as described above, as well as separately normalized using RPKM (reads per kilobase per million) when within-sample comparison was necessary. Prokaryotic viral diversity was calculated using the Shannon-Wiener index as part of the vegan R package (v2.6)(99). The mean index was generated by 1000 bootstraps and a t-test was done on the differences of means to determine significance.

Abundance (metagenomic) and activity (metatranscriptomic) of each virus were used to show the proportional abundance and activity of taxonomically categorized viruses at each zone.

### Eukaryotic Virus Diversity and Abundance

Viruses belonging to the phylum *Nucleocytoviricota* (NCLDV) were recovered from each zone’s metagenomic and metatranscriptomic dataset using the major capsid protein (MCP) marker gene. Briefly, proteins were predicted from each metagenome and metatranscriptome using prodigal-gv (v2.11)(100) and MCPs were recovered and identified using the HMM-based tool NCLDV-Markersearch (101). MCPs recovered were then subjected to a 150aa length cutoff before being dereplicated at 95% amino acid identity using cd-hit (102). This yielded a total of 33 MCP candidates from all zones, which were aligned to reference sequences from the GVDB (101) using MAFFT (v7.520)(103) with the ‘--auto’ setting. This alignment was trimmed using trimAl (v1.2)(104) with the ‘-gt 0.1’ parameter. IQ-TREE (v1.6.12))(105) was used to build a maximum-likelihood tree of the alignment using the ‘LG+F+R10’ model with 1000 bootstraps. The resulting tree was visualized using iTOL (106). Taxonomy was assigned to each MCP based on phylogenetic placement as well as with gvclass (107) which also uses a tree-building method. Abundance and activity were quantified by mapping metagenomic and metatranscriptomic reads to MCPs at 95% minimum identity with minimap2. Coverage was generated using CoverM and TMM normalization was performed on raw read mapping values

### Polintons, PLVs, Virophages

Polintons, Polinton-like Viruses, and Virophages are all united in their shared major capsid structure (108). Using this information, a set of custom HMM profiles specific to diverse capsid proteins in these viruses were created as reported in Stephens et al. (109). using HMMER3 (v3.4) (110) with an e-value cutoff of 1e-5. After applying a >200 amino acid length cutoff, MCPs were dereplicated at 95% identity using cd-hit resulting in 183 total MCPs. These MCPs were aligned with reference sequences from Stephens et al. (109) using MAFFT with the ‘--auto’ flag and the alignment was trimmed using trimAl with the ‘-gt 0.1’ option. IQ-TREE was used to build a maximum-likelihood tree of the alignment using the ‘LG+F+R10’ model with 1000 bootstraps. The resulting tree was visualized using iTOL. The same read mapping protocol and normalization as stated above were used to quantify the abundance and activity of these elements.

### RNA virus diversity and abundance

RNA viruses were identified in the metatranscriptomic datasets using the RdRp-scan pipeline (111). Proteins were predicted from assembled contigs using prodigal (v2.6.3)(112) and proteins were dereplicated at 98% identity using cd-hit. These proteins were then scanned using HMMER3 with the RdRp hmm profile. Recovered RdRp candidates were only kept if they were >200aa and were then screened for A, B, and C motifs using the RdRp motifs database (111). The resulting RdRp candidates were aligned to reference sequences from the known RNA virus database (111). This alignment was subject to trimming using trimAl with ‘-gt 0.1’ parameter. IQ-TREE was then used to make a phylogenetic tree of the alignment using the ‘LG+F+R10’ model with 1000 bootstraps. In total, 143 RdRp sequences were recovered. Metatranscriptomic reads were mapped to the RdRp genes to get abundance information. Read mapping and normalization procedure was the same as used for NCLDVs and other eukaryotic viral elements.

### Host prediction of prokaryotic viruses

As part of the ViWrap pipeline, iPHOP predicts hosts of prokaryotic viruses using a combination of host-based and virus-based methods (113). A total of 36 virus-host linkages were predicted with a confidence score of over 90 using these methods. The resulting virus-host linkages were visualized using the Circos R package (114).

Additional hosts were predicted using CRISPR spacers inside of bacterial bins obtained from the datasets. Briefly, assembled contigs from each zone were binned using metabat2 (115), and resulting bins were classified with the GTDB-TK classify tool (v1.6.0)(116). CRISPR spacers were identified in the bacterial bins using the web based CRISPR Recognition Tool (CRT) (v1.1) (117) and then SpacePHARER (v5)(118) was used to match these CRISPR spacers to recovered virus MAGs. This method yielded a total of 9 additional host predictions using an e-value cutoff of 1e-5.

### Active viral ecology and gene expression

After mapping reads to prokaryotic virus MAGs, these viruses were categorized based on their activity and abundance across zones. To elucidate trends in activity and abundance, viruses were assigned abundance and activity rankings for each zone. These rankings were used to first establish a “low-abundance, non-active” cutoff based on the point where the rank abundance curve flatlines for both abundance and activity. The remaining viruses were assigned an “Activity: Abundance” ratio that was used to cluster viruses by Euclidean distance. This hierarchical clustering yielded 3 main groups which were classified by investigating the average ratios of activity to abundance (RPKM). Based on the activity: abundance ratios in these three groups, viruses were separated into “Highly active” – viruses with very high activity:abundance ratios; “Active + Abundant” – viruses with activity:abundance ratios near 1; and “Abundant, non-active” – viruses with activity:abundance ratios less than 1.

Based on clustering, virus MAGs were separated into two groups to analyze differences in gene expression. A total of 178 “highly active” viruses were identified and these were then compared to the 1006 viruses from other groups. Genes and proteins were predicted for both groups of viruses using prodigal. Proteins were annotated using the Pfam (119), VOG (120), and eggNOG (121) databases with HMMER3 using an e-value cutoff of 1e-5. Metatranscriptomic reads were mapped to genes using CoverM (contig mode) at 95% minimum identity. Raw read counts were normalized using RPKM to normalize for gene length and total reads mapped. Grouped by annotations, gene expression from all zones was summed together to get comprehensive expression across zones.

To analyze differences in gene expression between zones, viruses were grouped into zone-specific expression groups based on their TMM-normalized total expression at one zone compared to the others. A virus was deemed to be “zone-specific” if its total expression at one zone was greater than 10× higher than the other two zones combined and it had a TMM value of greater than 1. The resulting groups were plotted in a 3D scatter plot using the scatterplot3d R package (v0.3-44)(122). Once these zone-specific groups were established, the top 18 highly expressed genes in each of those groups were obtained and visualized using the pheatmap R package (v1.0.12)(123).

Auxiliary metabolic genes (AMGs) were obtained from VIBRANT (95) output as part of the ViWrap pipeline. Reads were mapped to AMGs separately at 95% identity with CoverM (contig mode). Read counts were normalized using RPKM and plotted as z-scores using pheatmap. Genes annotated as phage integrases were also separated to compare integrase expression across zones. TMM values for the sum integrase expression for each zone was normalized by dividing by the total TMM viral expression for that zone to account for there being unequal total viral expression across zones.

### Jumbo phage identification and host analysis

Two jumbo phages were identified from the ViWrap outputs based on genome size (>200 kbp). Each of these jumbo phage genomes was annotated and plotted using pharokka (124). Both jumbo phages were linked to host bacterial bins using CRISPR spacers and these bins were subject to metabolic investigation.

For manganese oxidation, the mnxG gene expression was used as a proxy for the entire process (125). A custom hmm profile for mnxG was created using reference sequences of bacterial mnxG from the UniProt database (126). This profile was used to search for mnxG genes in all bacterial bins using HMMER3 with an e-value cutoff of 1e-5. A similar method was used to recover the arsM genes for arsenic metabolism. After gene recovery, reads were mapped to these genes at 95% minimum identity using minimap2 and CoverM contig mode. Reads were normalized using the TMM method and then plotted as heatmaps using z-score normalization of rows for within-zone comparisons.

Reads were also mapped to the entire jumbo host bacterial bins at 95% to look at host gene expression patterns. These read counts were normalized using RPKM and grouped into KEGG pathways (127) based on annotations from the eggNOG database.

## Data Availability

Raw reads are available on NCBI Bioproject PRJXXXXX. Other data such as phylogenetic tree files, alignments, protein annotations, phage genomes, and genome statistics are available on Figshare (https://figshare.com/projects/Active_prokaryotic_and_eukaryotic_viral_ecology_in_a_deep-sea_brine_pool/191886). The code used to generate graphs and analyze data is also available on Figshare.

## Acknowledgements

We owe a debt of gratitude to our Saudi Arabian partners, especially NEOM, for supporting this study. We are similarly indebted to OceanX and the crew of OceanXplorer for their operational and logistical support for the duration of this expedition. In particular, we would like to acknowledge Andrew Craig, Olaf Dieckoff, Ewan Bason, Larissa Frühe, and Kate von Krusenstiern for data acquisition, sample collection, and support of scientific operations aboard OceanXplorer. We would also like to thank OceanX Media for documenting and communicating this work with the public. SP and MC were funded by NEOM under Agreement No: SRA-ENV-2023-001 / AWD-008854.

## Notes

https://figshare.com/projects/Active_prokaryotic_and_eukaryotic_viral_ecology_in_a_deep-sea_brine_pool/191886

### Competing Interest Statement

The authors have declared no competing interest.

