## Supplemental Figures for "Active prokaryotic and eukaryotic viral ecology across spatial scale in a deep-sea brine pool": Brine Pool Supplemental.docx

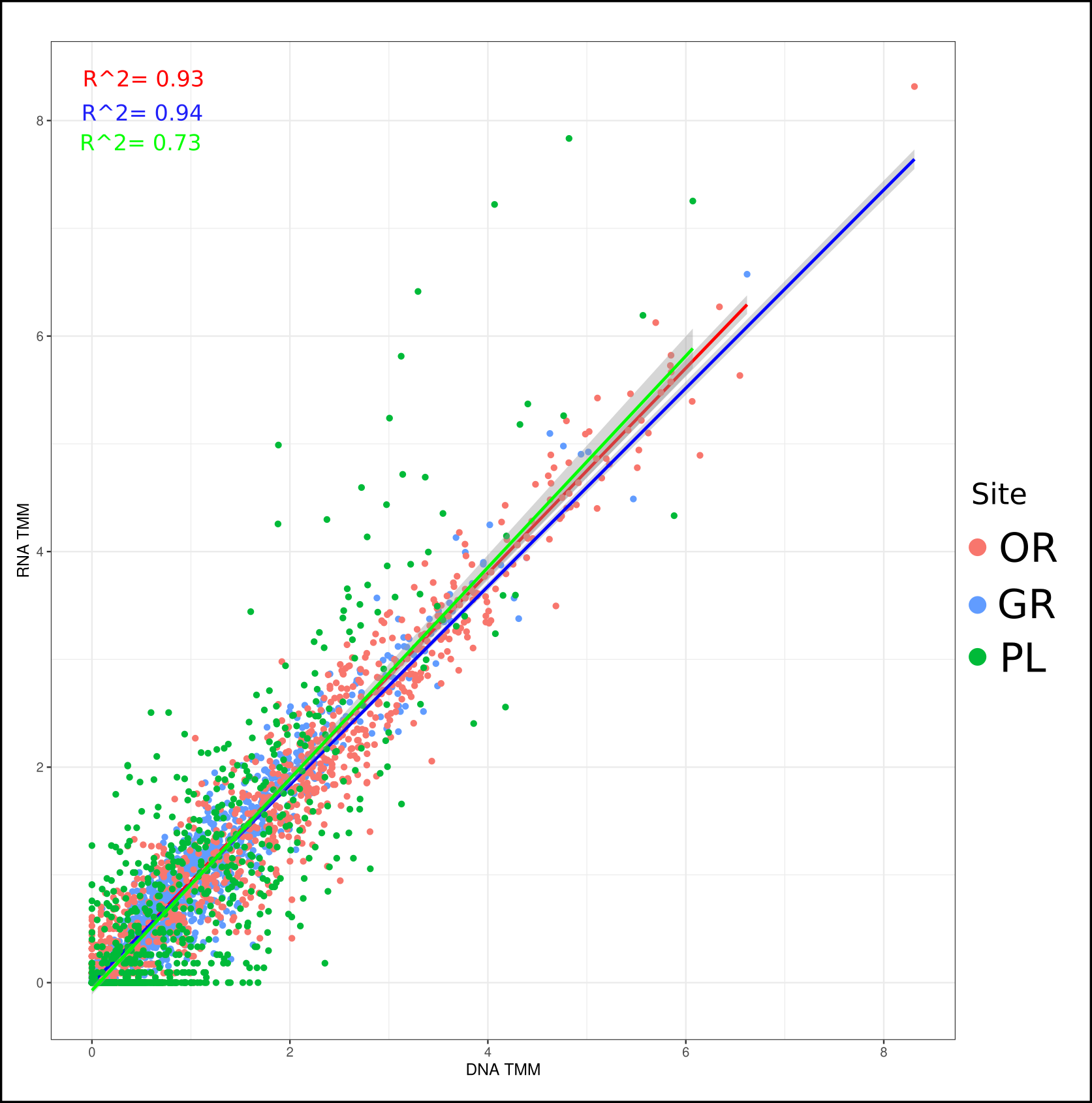

**Figure S1. Relationship between abundance and activity of the virus community.** Metagenomic (DNA) and metatranscriptomic (RNA) data for each virus at each zone was plotted on a scatter plot, with each point representing a single virus. Regression lines were drawn with a standard linear regression and R^2 values are displayed.

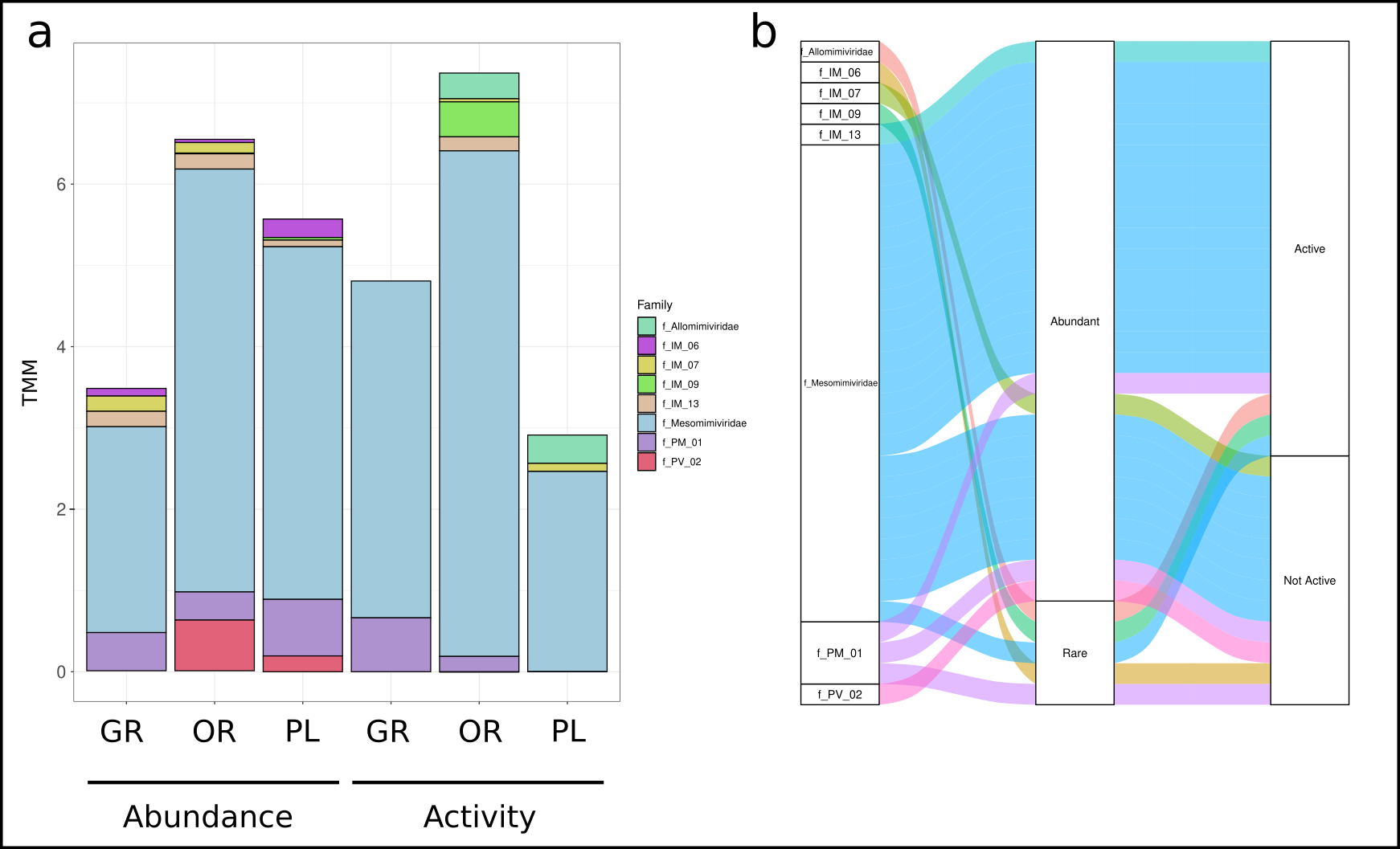

**Figure S2. Giant virus (NCLDV) abundance and categorization. (a)** TMM normalized abundance and activity of each NCLDV MCP is displayed here. Family-level classifications were determined putatively using GV-class. **(b)** An alluvial plot showing all identified NCLDV families as well as their patterns of abundance and activity. An NCLDV was determined to be abundant if it had a read count of > 5 and active if it had a similar cutoff of metatranscriptomic reads.

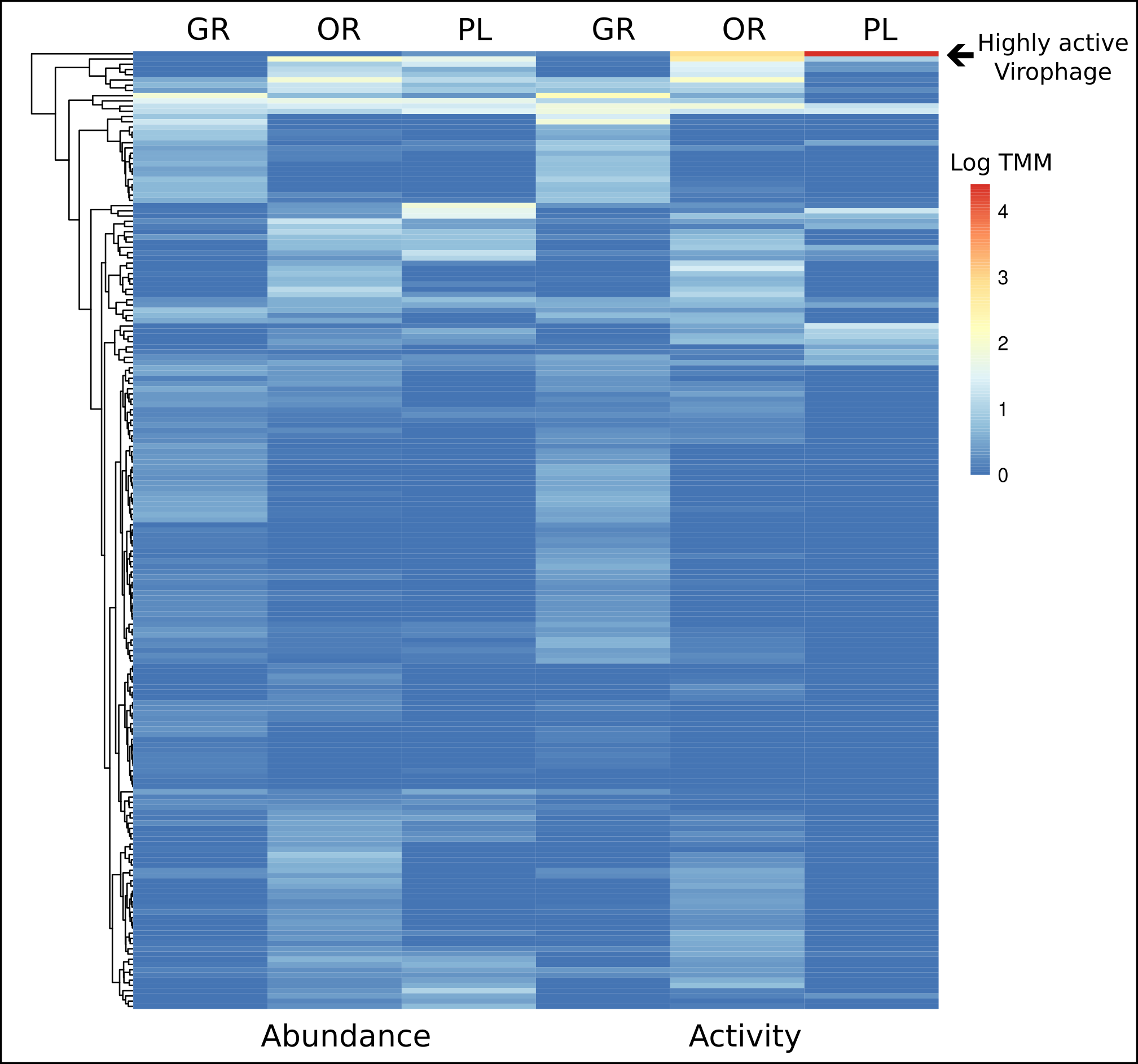

**Figure S3. Virophage activity across zones.** Abundance and activity data for each virophage is displayed on this heatmap using log TMM values. The highly active virophage is highlighted at the top of the graph.

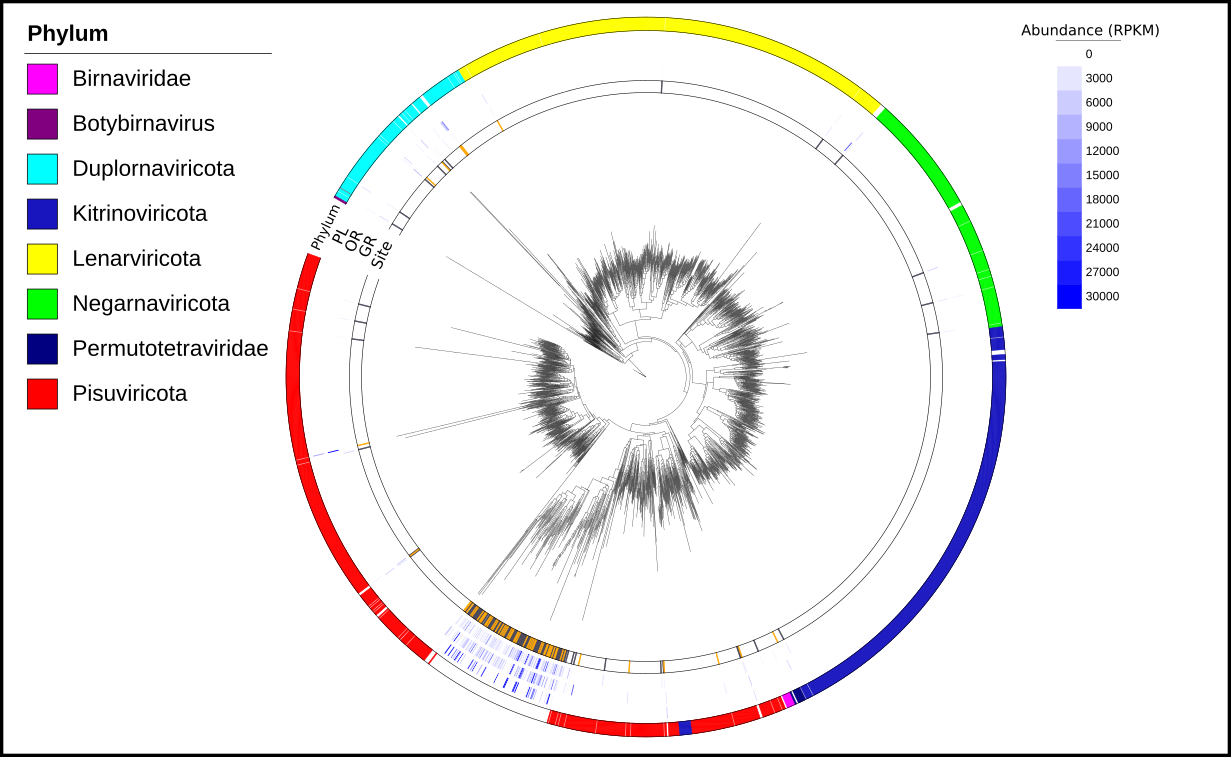

**Figure S4. RNA virus diversity and abundance across zones**. A phylogenetic tree displaying RdRp sequences from 8 major RNA virus phyla using reference sequences from the RdRp scan database. Recovered RNA viruses from each zone at the brine pool are shown in orange and gray bands on the “site” track of the tree. Metatranscriptomic reads were mapped to recovered viruses to get abundance at each zone, normalized using RPKM.

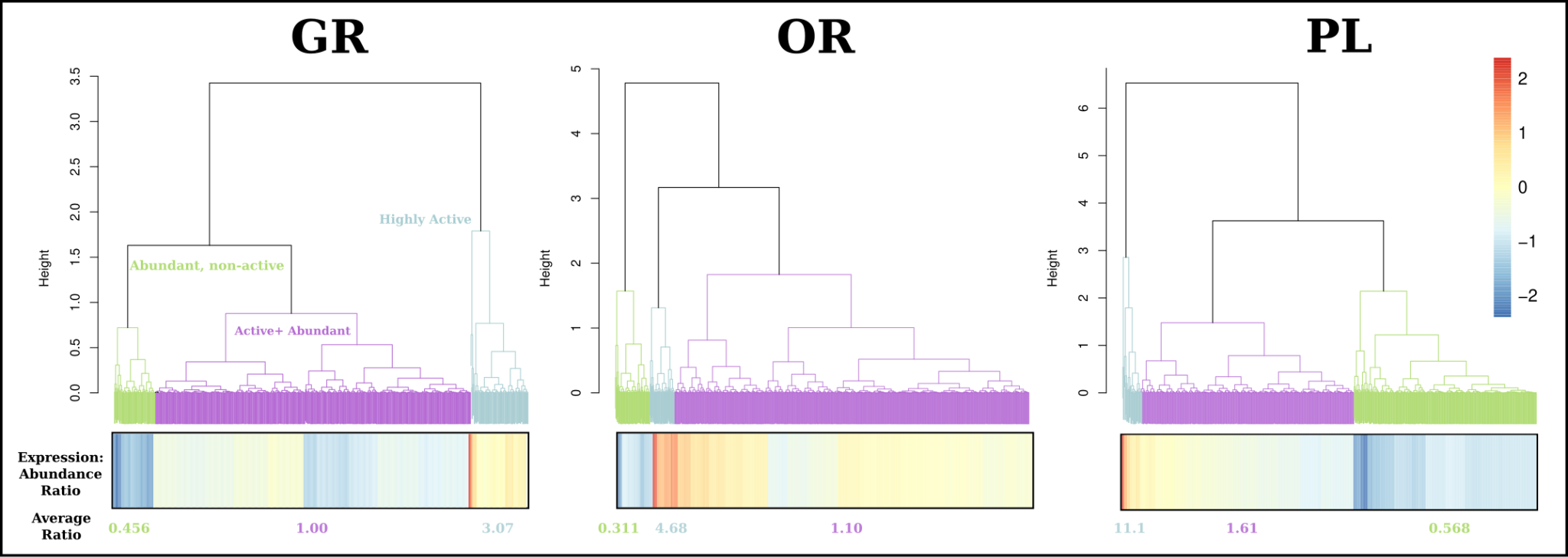

**Figure S5. Hierarchical clustering of active viruses.** Hierarchical clustering based on a distance matrix created from expression:abundance ratios was performed for each zone. A heatmap of this ratio was also created and displayed under the tree to see the patterns. Average ratio of each group was calculated as well and used to confirm clusters.

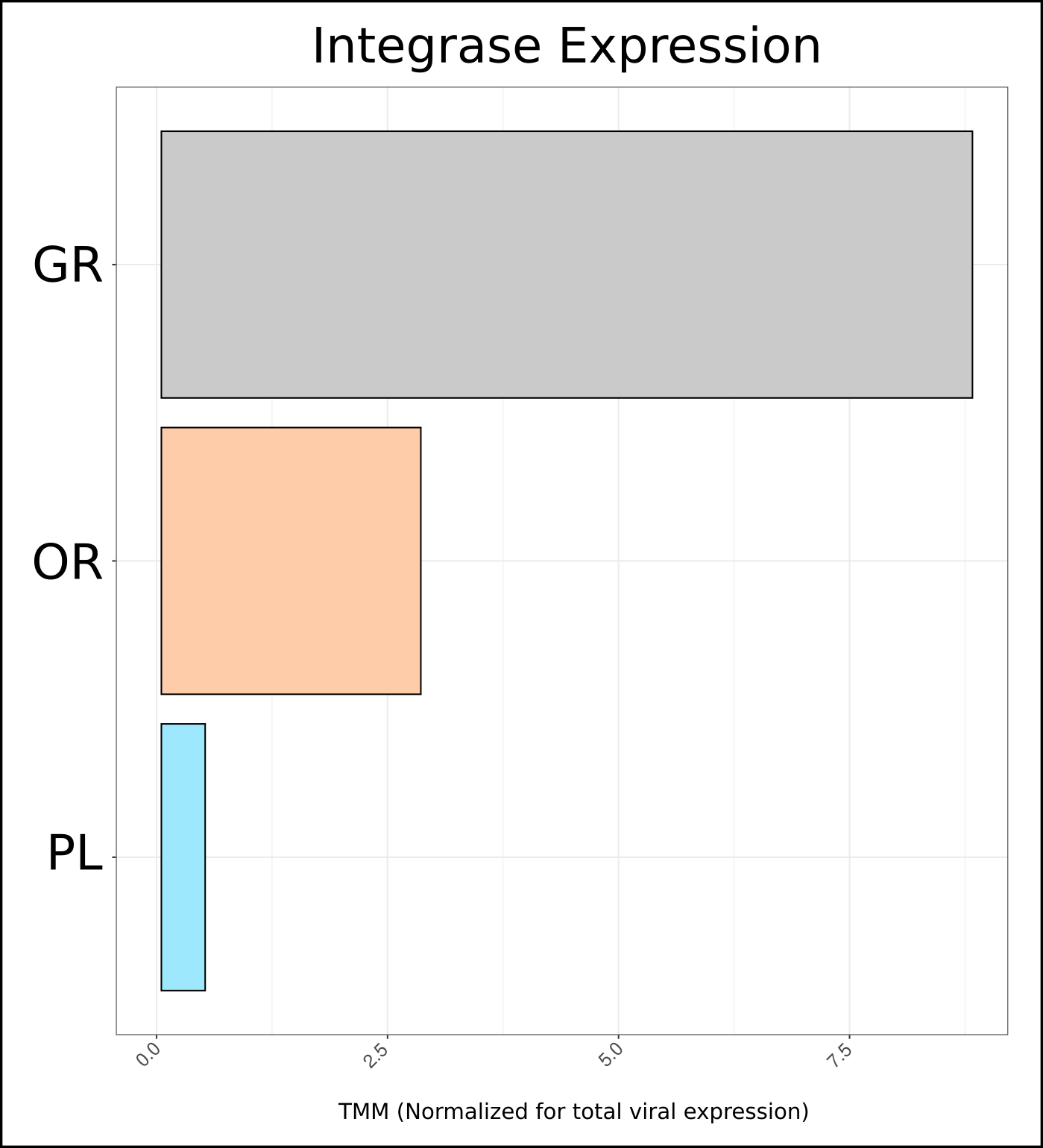

**Figure S6. Integrase expressions across zones.** Integrase genes from each zone were retrieved from protein annotations and total expression was summed together. This total expression was then normalized for total viral expression at each zone to account for there being more viruses at the gray zone compared to the other two.

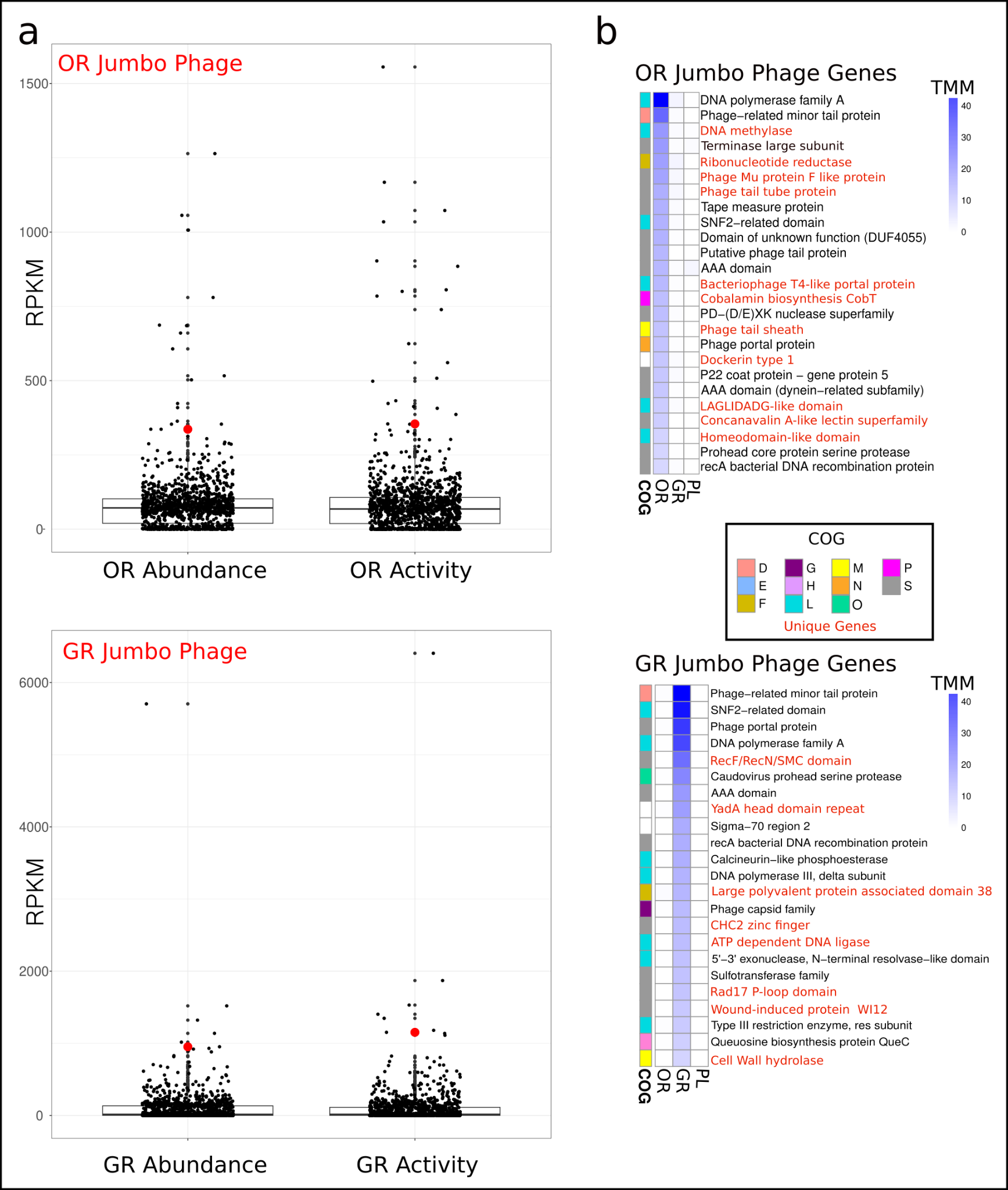

**Figure S7. Jumbo phage within-zone ecology and gene expression.** **(a)** Placement of OR and GR jumbo phages within the larger viral community at their respective zones. Each dot represents either the abundance or expression of a single virus, while the red dots represent the jumbo phages. **(b)** Gene expression for each jumbo phage was done by mapping reads to their genomes. The top 25 most expressed genes are displayed here with red genes representing genes present in one jumbo phage but absent in the other. COG categories were also assigned based on eggNOG annotations [D: Cell cycle control, E: Amino acid metabolism, F: Nucleotide metabolism, G: Carbohydrate metabolism, H: Coenzyme metabolism, L: Replication and repair, M: Cell wall/membrane biogenesis, N: Cell motility, O: Post-translational modification, P: Inorganic ion transport, S: unknown].

| **Sample** | **Raw Reads** | **Trimmed Reads** | **Percent kept** | **Assembly length (bp)** |
| --- | --- | --- | --- | --- |
| GR_DNA | 119326035 | 119292693 | 0.999720581 | 1627118281 |
| OR_DNA | 109418220 | 109346733 | 0.999346663 | 1503273355 |
| PL_DNA | 12593395 | 12567966 | 0.997980767 | 124447755 |
| GR_RNA | 93858854 | 93834516 | 0.999740696 | 734505704 |
| OR_RNA | 108728598 | 108695355 | 0.999694257 | 352891040 |
| PL_RNA | 8059986 | 8054264 | 0.999290073 | 8928061 |

**Table S1. Trimming and assembly statistics.** A table containing the number of raw reads for each sample (grey zone: GR; orange zone: OR; pool zone: PL). Trimming and assembly were performed using trimGalore and MEGAHIT respectively.
